# A complete MAPK cascade, a calmodulin, and a protein phosphatase act downstream of CRK receptor kinases and regulate *Arabidopsis* innate immunity

**DOI:** 10.1101/2022.03.27.486008

**Authors:** Fangwen Bai, Johannes W. Stratmann, Daniel P. Matton

## Abstract

Mitogen-activated protein kinase (MAPK) cascades are critical signal transduction modules in stress responses, but how their composition and mode of activation induces a stress response is poorly understood. We showed in Arabidopsis that CRK21, a cysteine-rich receptor-like protein kinase (CRK), phosphorylates MAPK kinase kinase 20 (MKKK20) and thus directly activates a novel MAPK cascade, consisting of MKKK20, the MAPK kinase MKK3, and the MAPK MPK6. Furthermore, the protein phosphatase PP2C76 and the calmodulin CaM7 were identified as negative and positive modulators of the cascade, respectively. Loss-of-function in components of the MAPK cascade or in CaM7 led to susceptibility to the bacterial pathogen *Pseudomonas syringae* and the fungal pathogen *Botrytis cinerea*. In contrast, loss-of-function of PP2C76 as well as transient overexpression of the genes in the MAPK cascade and CaM7 conferred resistance to the pathogens. Moreover, seven additional CRKs interacted with MKKK20 *in vivo*, and four of these were highly expressed after inoculation with *P. syringae*. In summary, our findings demonstrate that the novel CRK21-MKKK20-MKK3-MPK6 signaling pathway functions in immunity to fungal and bacterial pathogens and that CRKs may function in directly activating MKKKs.

## Introduction

Phytopathogens are one of the major threats to agriculture (Savary et al., 2019). They can be classified based on their lifestyle (Glazebrook, 2005). Necrotrophic pathogens like *Botrytis cinerea* kill their host cells rapidly after infection by secreting toxins, and they derive nutrients from dying or dead plant tissues. In contrast, biotrophic pathogens keep their hosts viable, and they obtain nutrients from living cells. Hemibiotrophic pathogens exhibit both biotrophic and necrotrophic characteristics. The bacterial pathogen *Pseudomonas syringae* has a biotrophic lifestyle in the early stage of infection but eventually kills its host plant (Glazebrook, 2005; Panthapulakkal Narayanan et al., 2020).

Pathogen or microbe-associated molecular patterns (PAMPs/MAMPs) are derived from pathogens and perceived by membrane-bound pattern recognition receptors (PRRs), which leads to PAMP-triggered immunity (PTI) (Bigeard et al., 2015). PTI can be overcome by plant pathogens via effector proteins that are secreted into plant cells to promote pathogenesis. The effector proteins can be detected and recognized by cytosolic or extracellular receptors encoded by resistance genes, which results in effector-triggered immunity (ETI) to suppress pathogenesis (Cui et al., 2015).

The PRR-mediated recognition of PAMPs is a key step in the initiation of an effective plant defense response (Zipfel, 2008). Plant PRRs are either receptor-like kinases (RLKs) or receptor-like proteins (RLPs), both of which contain a ligand-binding extracellular domain and a transmembrane domain. RLKs also possess a cytosolic kinase domain (Boutrot and Zipfel, 2017). Upon binding of PAMPs to PRRs, RLKs will undergo homo-dimerization or heterodimerize with membrane-localized coreceptors to form a receptor complex, resulting in signal initiation through phosphorylation (Chinchilla et al., 2009; Hohmann et al., 2017).

The leucine rich repeat receptor kinases (LRR-RKs) FLAGELLIN SENSING2 (FLS2) and EF-TU RECEPTOR (EFR) are two well-studied PRRs in Arabidopsis which function in the recognition of the bacterial PAMPs flg22 (a 22-amino-acid epitope of bacterial flagellin) and elf18/elf26 (an N-acetylated 18/26 amino-acid epitope of bacterial elongation factor Tu), respectively (Chinchilla et al., 2006; Zipfel et al., 2006). Once the corresponding PAMPs are perceived by FLS2 and EFR, the RLKs heterodimerize with the coreceptor BRASSINOSTEROID INSENSITIVE1-ASSOCIATED KINASE1 (BAK1) to form active receptor complexes and consequently initiate plant defense (Roux et al., 2011; Sun et al., 2013; Tang et al., 2015). The membrane-localized protein BOTRYTIS-INDUCED KINASE1 (BIK1), involved in the resistance to necrotrophic and biotrophic pathogens in Arabidopsis, is transcriptionally regulated by *B. cinerea* infection (Veronese et al., 2006). As a receptor-like cytoplasmic kinase (RLCK), BIK1 is a direct substrate of PRR-coreceptor complexes and plays a key role in integrating signals from PRRs like FLS2, EFR, or LYK5. In addition, these PRRs activate mitogen-activated protein kinase (MAPK) cascades that are crucial in plant resistance to pathogens attack (Bi et al., 2018).

Cysteine-rich receptor-like kinases (CRKs) are a subfamily of RLKs with more than 40 members in Arabidopsis. They are characterized by the presence of two DUF26 (Domain of Unknown Function 26) domains in the extracellular part of the receptor, which contain four conserved cysteines. A number of CRKs function in plant immunity. The expression of CRK4, 5, 6, 10, 11, 19, and 20 is induced by pathogen infection. CRK4, 5, 19, and 20 also trigger rapid cell death (Chen et al., 2003; Chen et al., 2004a). Increased expression of CRK4, CRK6, CRK13, CRK28, CRK36, and CRK45 triggers PTI-like responses and enhances the resistance of Arabidopsis to *P. syringae*, and *crk45* mutant plants are more sensitive to virulent bacteria (Acharya et al., 2007; Yadeta et al., 2017; Zhang et al., 2013). CRK4, CRK6, and CRK36 interact with FLS2. CRK36 also associates with BIK1 further enhancing BIK1 phosphorylation (Lee et al., 2017; Yeh et al., 2015) and functioning as a negative regulator in the response to infection by *Alternaria brassicicola*, a necrotrophic fungal pathogen.

MAPK signaling cascades are universal signal transduction modules, regulating a wide range of biological processes in eukaryotes (Bigeard and Hirt, 2018; He et al., 2020; Kumar et al., 2020; Zhang et al., 2018). The classical three-tiered MAPK cascade comprises a MAPK kinase kinase (MAPKKK, MKKK or MEKK) that phosphorylates and activates a MAPK kinase (MAPKK, MKK or MEK), which in turn phosphorylates and activates a MAPK (MPK). Finally, the activated MPK will phosphorylate downstream substrates such as transcription factors to regulate growth, development, and responses to biotic and abiotic stresses (Lee et al., 2015). Several MPKs function in the plant immune response. MPK3/6, MPK9 and MPK12 are essential for stomatal immunity (Salam et al., 2013; Su et al., 2017), and MPK3, MPK4, MPK6, MPK9, and MPK12 are all involved in the response to *P. syringae* infection (Desikan et al., 2001; Jammes et al., 2011; Nie and Xu, 2016). MPK3 and MPK6 also function in the resistance to *B. cinerea* infection (Han et al., 2010). Nevertheless, to date, only few complete MAPK cascades have been identified in Arabidopsis immunity, such as the MEKK1-MKK1/2-MPK4 cascade, which negatively regulates immunity (Gao et al., 2008; Kong et al., 2012; Suarez-Rodriguez et al., 2007; Takagi et al., 2019), and the MEKK1-MKK4/5-MPK3/6 (Asai et al., 2002), which positively regulates immunity, although both cascades are induced by flg22.

Once the immune response is initiated, the intensity and duration of MAPK activation must be tightly regulated to obtain a specific signaling outcome. Protein phosphatases dephosphorylate MAPKs and are the main negative regulators of MAPK signaling pathways. MAPK phosphatases (MKPs) include the type 2C protein phosphatases AP2C1, AP2C2, AP2C3, and AP2C4 of the PP2C clade B (Schweighofer et al., 2004), which directly inactivate MPK3, MPK4, and MPK6 (Brock et al., 2010b; Schweighofer et al., 2007; Umbrasaite et al., 2010). *ap*2*c1/3* double mutant plants show enhanced activation of ABA-induced MPK3 and MPK6 (Brock et al., 2010a), and *ap*2*c*1 null plants are less sensitive to *P. syringae* DC3000 infection (Shubchynskyy et al., 2017). In addition, HAI1, HAI2, and HAI3 of the PP2C clade A (Schweighofer *et al*., 2004) inactivate MPK3 and MPK6 and are activated by *P. syringae* DC3000 to suppress immunity (Mine et al., 2017).

Some plant MKPs are regulated by calmodulins (CaMs). CaMs are calcium (Ca^2+^) sensors, each containing a long and flexible linker that connect two independently-folded lobes, which make it a perfect adaptor or bridging protein (Villalobo et al., 2017). CaMs can modulate the activation of MKPs or directly act upon MAPK cascades (Yamakawa et al., 2004). In Arabidopsis, the phosphatase activity of MKP1 is enhanced by AtCaM2 in a Ca^2+^-dependent manner (Lee et al., 2008a). Conversely, the activity of the Arabidopsis MKP DsPTP1 is inhibited by Ca^2+^-activated CaM resulting in increased MPK6 activity (Kim et al., 2021). In Arabidopsis, CaM3, CaM4, and CaM7 activate MPK8 in concert with MKK3 in wounding responses (Takahashi et al., 2011), and in response to cold, cytosolic calcium spikes stimulate the binding between CaM and Calcium/Calmodulin-regulated Receptor-like Kinase1 (CRLK1) which activates the MEKK1-MKK2-MPK4/6 cascade (Furuya et al., 2013; Yang et al., 2010a; Yang et al., 2010b; Yuan et al., 2018).

Arabidopsis MKKK20 belongs to the MEKK-like kinases of subclade III (Colcombet et al., 2016). It was shown to interact with CaM and CaM-like proteins (CML) in protein microarrays (Popescu et al., 2007). In the osmotic stress response, MKKK20 activates MPK6 (Kim et al., 2012). *mkkk20* mutants were hypersensitive to osmotic stress and exhibited weaker MPK6 activity after NaCl, drought, cold, and H_2_O_2_ treatments. In contrast, MKKK20 overexpressing plants showed resistance to salt stress (Kim *et al*., 2012). In the response to ABA, MKKK20 is part of the MKK5-MPK6 cascade which regulates primary root growth and the stomatal response (Li et al., 2017).

Earlier, we showed that the two separate signaling pathways MKKK20-MKK3 and MKKK20-MPK18 are involved in microtubule functions (Benhamman et al., 2017). In the present work, we investigated the role of Arabidopsis MKKK20 in plant immunity. Here, we demonstrate that MKKK20 forms a novel MAPK cascade with MKK3 and MPK6, and this MAPK cascade is located downstream of CRK21 and other CRKs. We also show that the signaling module MKKK20-MKK3-MPK6 is negatively influenced by the protein phosphatase PP2C76 and positively influenced by the calmodulin CaM7, and that it functions in the immune response to plant pathogens. This report provides an unprecedented level of detail of a novel signaling pathway, including a RLK which directly activates a MAPK cascade via phosphorylation of a MKKK, and a positive (CaM7) and a negative modulator (PP2C76) of the cascade.

## Results

### MKKK20 interaction patterns

Recently, we reported that MKKK20 interacts with MKK3 to regulate cortical microtubule function, and that MKKK20 can also directly interact with MPK18 (Benhamman *et al*., 2017). However, no functional upstream or downstream factors were identified. Based on a large-scale yeast two-hybrid (Y2H) assay, numerous MKKK20-interacting candidate proteins had been identified (Benhamman *et al*., 2017). To further investigate the connections among the candidates from this Y2H screen, we analyzed them along with MKKK20 in the Arabidopsis Network Analysis Pipeline (ANAP) (http://gmdd.shgmo.org/Computational-Biology/ANAP) that integrates numerous protein interaction databases and reveals potential protein interaction networks with both direct and indirect interactions (Wang et al., 2012). After elimination of indirectly connected proteins, we retained a calcium-binding EF-hand family protein, numerous calmodulins (CaM), and a calmodulin-like protein (CML) that are known to play key roles in regulation of MAPK signaling pathways. The receptor-like kinase CRK21 was found to interact with CaM1 and CaM4 which also interact with MKKK20 (Figure 1A). We had shown previously that CRK21 directly interacts with MKKK20 in a Y2H screen (Benhamman *et al*., 2017). We verified these interactions in a directed Y2H assay (Figure 1B) and by BiFC (Bimolecular Fluorescence Complementation) (Figure 1C). The resolution of our BiFC assay is insufficient to draw conclusions about the precise subcellular localization of the interactions.

**Figure 1.**
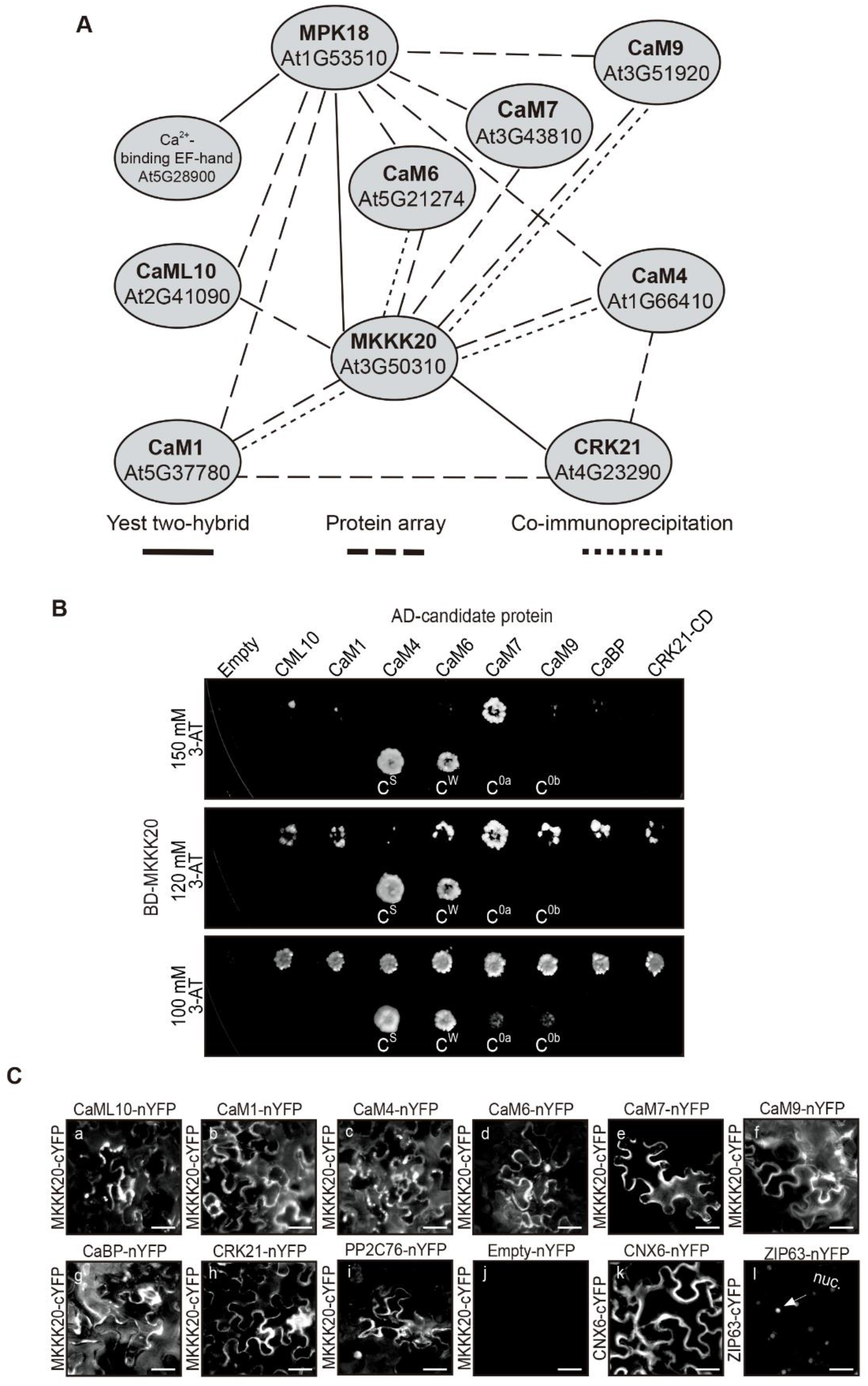
MKKK20 interaction network. (**A**) Network of MKKK20 interacting proteins from ANAP analysis and our previous Y2H screen. Lines between proteins represent the corresponding experimental methods that were used. (**B**) Validation of candidate genes from the ANAP analysis as MKKK20 interactors by a direct Y2H interaction assay. MKKK20 was fused to the GAL4 DNA-binding domain (BD), candidate genes to the GAL4-activating domain (AD), and both were transformed into yeast. Interaction strength was estimated using the following controls: C^s^: strong positive interaction control; C^w^: weak positive interaction control; C^0a^: negative interaction control; C^0b^: empty vector negative control. Images represent 1 out of 3 independent experiments. (**C**) Validation of candidate genes from the ANAP analysis as MKKK20 interactors by bimolecular fluorescence complementation (BiFC). MKKK20-cYFP was coexpressed with nYFP-tagged candidate interacting proteins in *N. benthamiana* epidermal cells through agroinfiltration. Protein–protein interaction was visualized by the fluorescence of YFP reconstitution. (a-i), BiFC assays demonstrating interactions between MKKK20 and candidate MKKK20-interacting proteins. (j), Coexpression of MKKK20-cYFP and nYFP (Empty) was used as a negative control. The self-binding cytoplasmic protein cofactor of nitrate reductase and xanthine dehydrogenase 6 (CNX6) (k) and nuclear protein ZIP63 (l) from Arabidopsis were used as positive controls. Images represent 1 out of 3 independent experiments. Scale bars, 50 μm.

### CRK21 directly activates MKKK20 and indirectly MKK3 *in vitro*

To confirm that CRK21 acts upstream of MKKK20, His-tagged CRK21 cytosolic domain (CRK21-CD) as well as MKKK20 and MKK3 were expressed in *E. coli* for in-solution kinase assays. As reported earlier (Benhamman *et al*., 2017), MKKK20 alone autophosphorylates, showing two major MKKK20 isoforms (Figure 2A; lanes 1, 3 and 4), while CRK21-CD alone did not autophosphorylate (Figure 2A; lane 2). However, phosphorylation of MKKK20 was strongly increased in the presence of CRK21-CD (Figure 2A; lane 3), suggesting that CRK21-CD either phosphorylated MKKK20, or increased autophosphorylation of MKKK20. To address this, CRK21-CD or MKKK20 were heat-denatured before adding them to the kinase reaction. Co-incubation of denatured CRK21-CD (d-CRK21-CD) and MKKK20 showed a similar level of MKKK20 phosphorylation as MKKK20 autophosphorylation (Figure 2A; lanes 4 and 1, respectively) while phosphorylation of d-MKKK20 was increased when incubated with CRK21-CD (Figure 2A; lane 5) compared to MKKK20 alone (Figure 2A; lane 1) and d-MKKK20 alone (Figure 2A; lane 7). Thus, presence of CRK21-CD strongly increased the phosphorylation of both MKKK20 and d-MKKK20.

**Figure 2.**
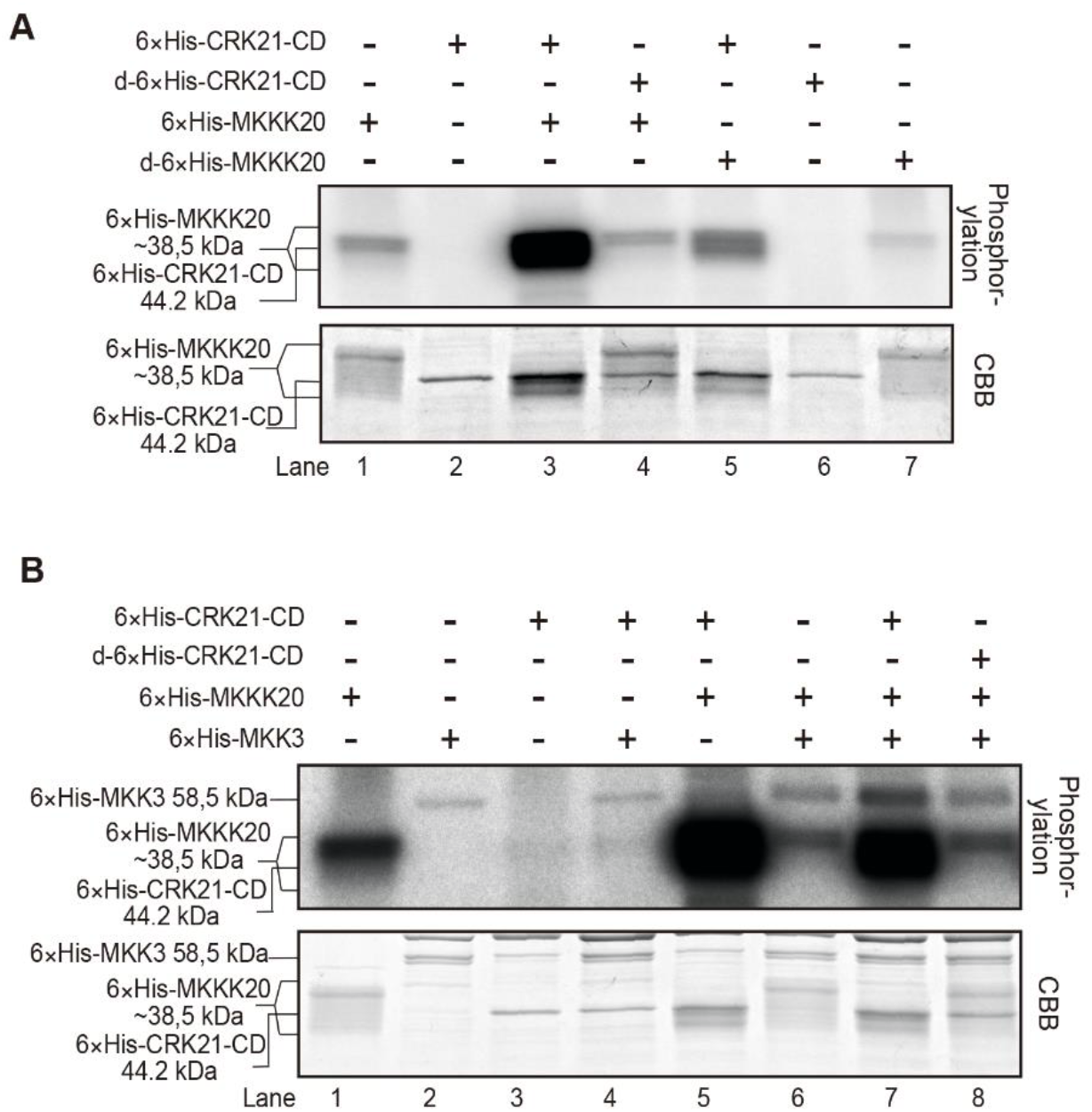
*In vitro* activation of MKKK20 and MKK3 by CRK21. (**A**) MKKK20 is phosphorylated by CRK21-CD. Phosphorylation signals (top) and CBB stained proteins (bottom) are shown. Note that purified His-tagged MKKK20 is not resolved as a narrow band, but rather a wide band based on phosphorylation status (compare lanes 1 and 7 to 3 and 5), unlike GST-tagged MKKK20 (Figure 5A and 5D). The wide band was verified by mass spectrometry as MKKK20 (see text). (**B**) CRK21-CD phosphorylates MKKK20 and enhances MKK3 phosphorylation. Phosphorylation signals (top) and CBB stained proteins (bottom) are shown. Tagged proteins were purified from bacterial extracts and incubated in kinase buffer containing [γ-^32^P]-ATP and subsequently analyzed by SDS-PAGE. To denature a subset of the proteins, they were boiled for 5 min. CBB, Coomassie Brilliant Blue stain. Images represent 1 out of 4 independent experiments.

Previously, MKK3 was shown to act downstream of MKKK20 (Benhamman *et al*., 2017). To determine interactions between MKK3 and CRK21-CD, MKK3 was incubated with MKKK20, CRK21-CD and MKKK20, and d-CRK21-CD and MKKK20, respectively (Figure 2B; lane 6, 7 and 8). As expected, MKK3 showed weak autophosphorylation (Figure 2B; lanes 2 and 4), while MKK3 phosphorylation was significantly enhanced in the presence of CRK21-CD and MKKK20 (Figure 2B; lane 7) but not by d-CRK21-CD-MKKK20 (Figure 2B; lane 8), confirming that CRK21 is an upstream receptor kinase of MKKK20 and MKK3. Increased MKK3 phosphorylation was not found in the presence of CRK21-CD alone (Figure 2B; lane 4) and depended on MKKK20 (Figure 2B; lane 7), confirming that MKKK20 is the upstream kinase for MKK3 (Benhamman *et al*., 2017). To better distinguish MKKK20 from CRK21-CD, which have similar apparent molecular weights, a GST-tag was fused to MKKK20 (Figure 5A). The kinase assays with 6xHis-GST-MKKK20 showed MKKK20 autophosphorylation and phosphorylation by CRK21-CD resulting in an additional band above autophosphorylated MKKK20 (compare Figure 5A; lane 2 to lane 3). Unlike in Figure 2A, CRK21-CD also showed phosphorylation in the presence of GST-MKKK20 (Figure 5A; lane 3). Together, these results suggest that, at least *in vitro*, CRK21 activates MKKK20 via phosphorylation resulting in MKK3 phosphorylation by MKKK20.

**Figure 5.**
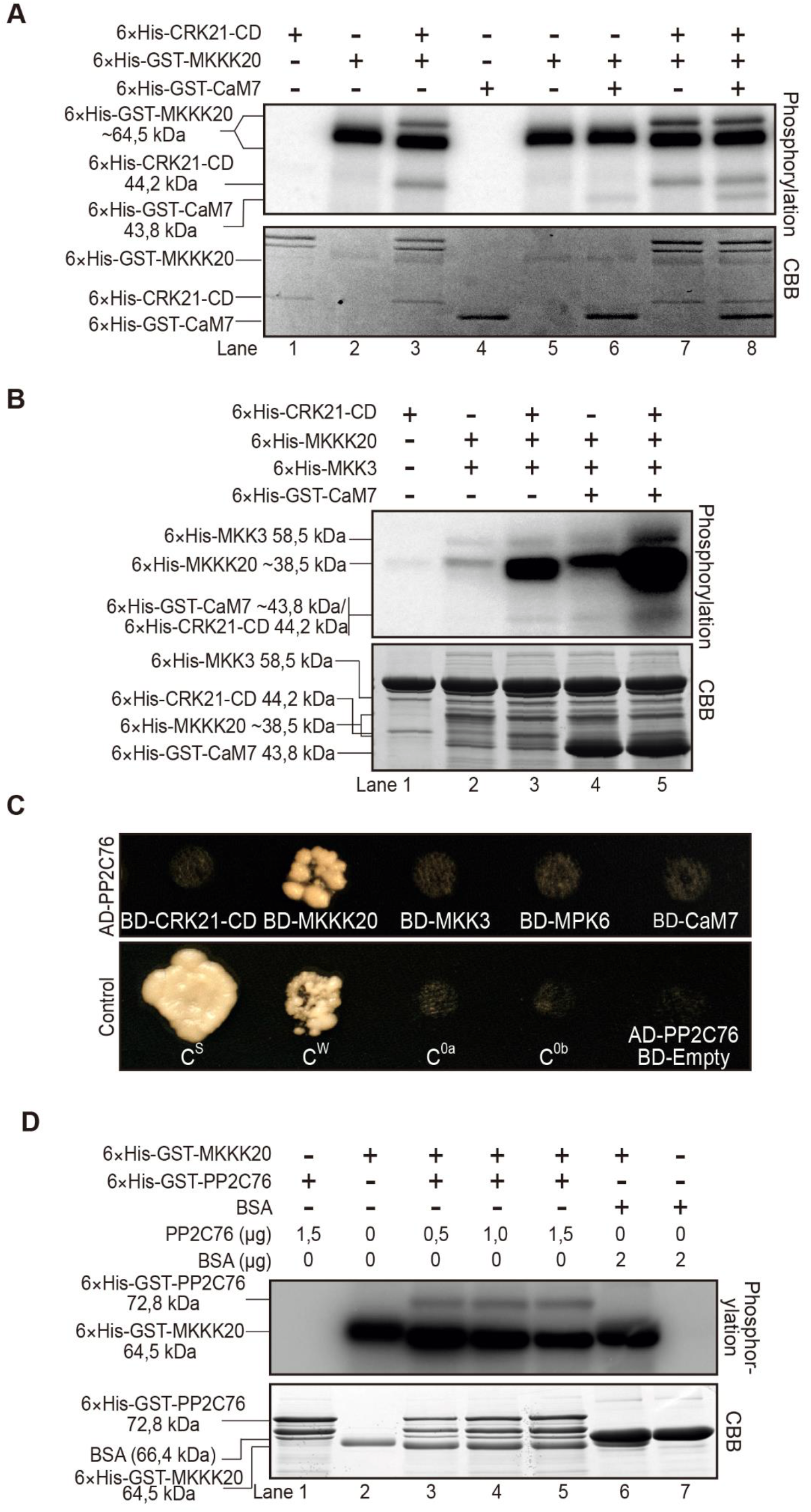
CaM7 and PP2C76 interact with the MKKK20 signaling cascade. (**A-B**) and (**D**), *In-vitro* kinase assays were performed as described in Figure 2. (**A**) CRK21-CD and CaM7 are phosphorylated by MKKK20. Phosphorylation (top) and CBB-stained proteins (bottom) are shown. (**B**) CaM7 enhances phosphorylation of CRK21-CD, MKKK20 and MKK3. (**C**) PP2C76 interacts with MKKK20. The directed Y2H assay was performed as describe in Figure 1B. PP2C76 was fused to AD while CRK21-CD, MKKK20, MKK3, MPK6, and CaM7 were fused to BD. Negative and positive controls are as shown in Figure 1B. (**D**) MKKK20 phosphorylates PP2C76, which dephosphorylates MKKK20. Phosphorylation (top) and CBB-stained proteins (bottom) are shown. See supplemental data Figure S1 for a related experiment. Note: expression and purification of recombinant proteins resulted in additional bands seen on CBB panels. Images represent 1 out of 3 (**A-C**) and 5 (**D**) independent experiments.

### Complex interactions of CRKs and MKKK20

To ascertain the potential roles of other CRKs in the CRK21-MKKK20-MKK3 signaling cascade, eight of the nine closest CRK paralogs of CRK21 were tested for their ability to also interact with MKKK20. They were selected based on their kinase domain phylogeny and include CRK12, CRK14, CRK16, CRK17, CRK18, CRK24, CRK30, CRK32, and the slightly more distant CRK33, which are all members of the CRK group IV (Bourdais et al., 2015). Six of the nine CRKs (CRK12, CRK14, CRK18, CRK24, CRK30, CRK33), strongly interacted with MKKK20 in Y2H and BiFC assays (Figure 3A and Figure 3B a-g). CRK16, which shares the most similar kinase domain with CRK21 and CRK24, showed a weak interaction with MKKK20 in a directed Y2H assay (Figure 3A), but an interaction in a BiFC assay comparable to other CRKs (Figure 3B; c). To further dissect the function of the CRKs, the cytosolic domains of CRKs that interacted with MKKK20 were expressed and used for autophosphorylation assays (Figure 3C) as well as for kinase assays with MKKK20 (Figure 3D). Among the CRKs, only CRK14-CD showed autophosphorylation (Figure 3C; upper and middle panel; lane 2). MKKK20 alone showed strong autophosphorylation (Figure 3D; lane 1). When combined, MKKK20 phosphorylated five of the seven CRK-CDs (CRKs 12, 14, 16, 18, and 24). The two bands for phosphorylated MKKK20 most likely show different phosphorylation patterns (Figure 3D) as shown by mass spectrometry (Benhamman *et al*., 2017) and by Li et al. (Li *et al*., 2017). Interestingly, the presence of CRK30-CD almost eliminated the MKKK20 autophosphorylation, suggesting that CRK30-CD may work as a negative regulator of MKKK20. Based on the strong autophosphorylation of MKKK20, it is difficult to determine MKKK20 phosphorylation by CRKs. It is possible that the extracellular domains of CRKs 30 and 33 are required for CRK kinase activity. While it is not clear whether other CRKs phosphorylate MKKK20, CRK21 clearly does (Figure 5A; lanes 2 and 3). This indicates a reciprocal phosphorylation between CRKs and MKKK20.

**Figure 3.**
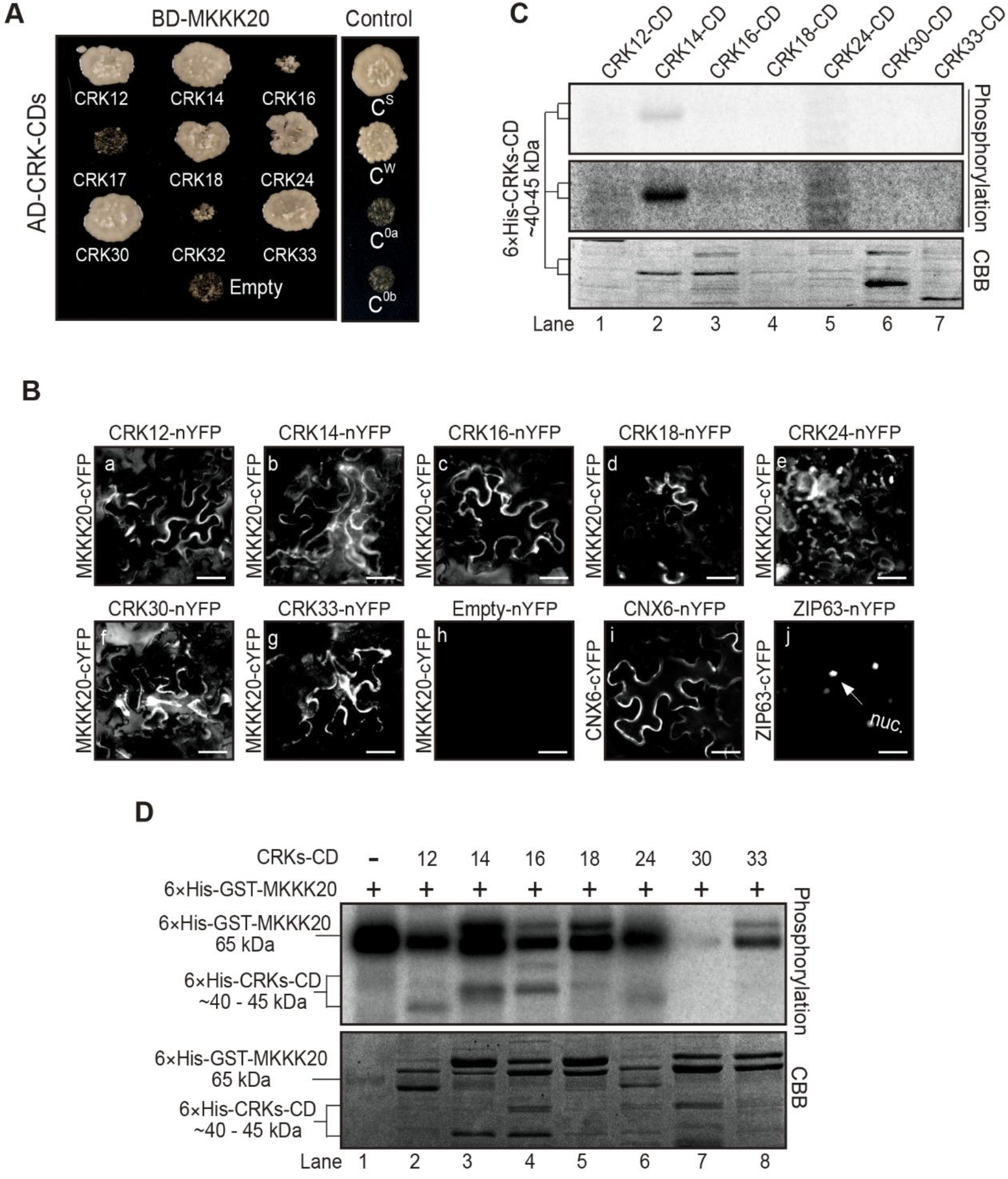
Several CRKs interact with MKKK20. (**A**) Y2H interaction between CRKs closely related to CRK21 and MKKK20. The directed Y2H assay was performed as described in Figure 1B. MKKK20 was fused to BD and the cytosolic kinase domains of CRKs (CRK-CDs) to AD. BD-MKKK20/AD-Empty served as a negative control. (**B**) Interaction of MKKK20 with various CRKs in BiFC assays. BiFC was performed as described in Figure 1C. MKKK20-cYFP was co-expressed with nYFP-tagged candidate-interacting proteins. Protein–protein interaction was visualized by the fluorescence of YFP reconstitution. BiFC assays showing interaction between MKKK20 and CRKs (a-g). Co-expression of MKKK20-cYFP and nYFP (Empty) was used as a negative control (h). CNX6 (i) and ZIP63 (j) were used as positive controls. Columns represent N-terminal YFP constructs and rows C-terminal MKKK20 constructs. Images in (**A**) and (**B**) represent 1 out of 3 independent experiments. Scale bars, 50 μm. (**C**) Autophosphorylation assay of MKKK20-interacting CRK-CDs. Top and middle panels show the same film with different exposure times. Images represent 1 out of 3 experiments. (**D**) Phosphorylation status of MKKK20 and MKKK20-interacting CRK-CDs. Kinase assays were performed as described in Figure 2. The image represents 1 out of 4 independent experiments.

### MPK6 functions downstream of CRK21-MKKK20-MKK3

MPK6 is activated by MKK3, MKK4, and MKK5 in response to JA and blue light (Sethi et al., 2014; Sozen et al., 2020; Takahashi et al., 2007). To determine if MPK6 could also act downstream of the CRK21-MKKK20-MKK3 cascade, the positive interaction between MKK3 and MPK6 was confirmed in Y2H and BiFC assays (Figure 4A and B). Moreover, the kinase dead version of MPK6 (MPK6-kd) was phosphorylated only when co-incubated with MKKK20 and MKK3 (Figure 4C; lane 5). To further confirm that MPK6 is the downstream target of CRK21-MKKK20-MKK3, the phosphorylation status of MPK6 was monitored *in vitro* in the presence of MKKK20/MKK3 or CRK21-CD/MKKK20/MKK3 (Figure 4D). MPK6 was weakly phosphorylated in the presence of MKKK20 and MKK3 (Figure 4D; lane 1) and strongly when CRK21-CD was added (Figure 4D; lane 2). The *in vivo* and *in vitro* data together suggest that MPK6 functions as the target of the CRK21-MKKK20-MKK3 cascade.

**Figure 4.**
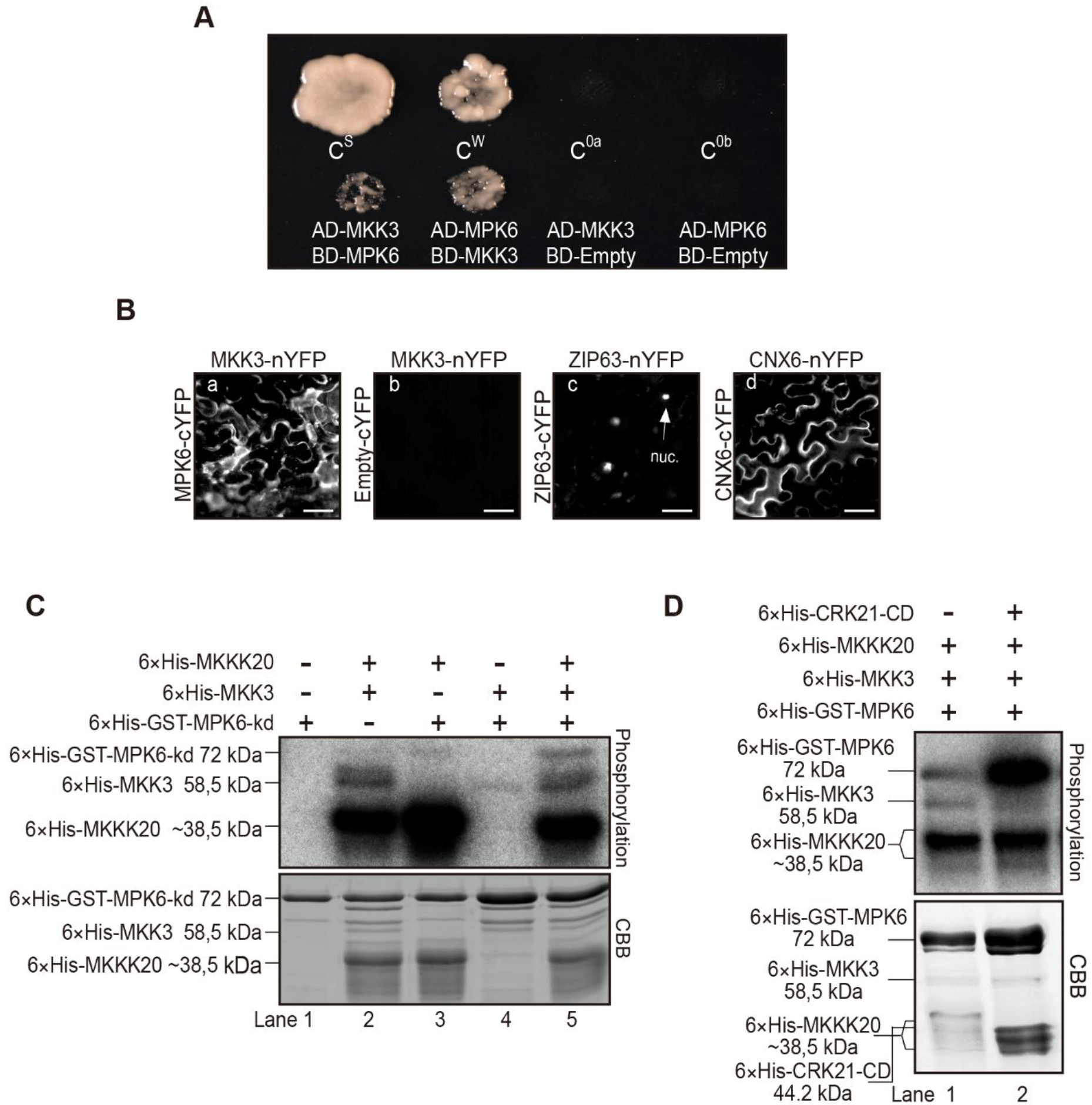
MPK6 functions downstream of the CRK21-MKKK20-MKK3 cascade. (**A-B**) MKK3 interacts with MPK6. (**A**) MKK3 interacts with MPK6 in a Y2H assay. The Y2H assay was performed as described in Figure 1B. MKK3 and MPK6 were fused to the GAL4 DNA-binding domain (BD) and GAL4-activating domain (AD), respectively. AD-MKK3/BD-Empty and AD-MPK6/BD-Empty served as negative controls. (**B**) MKK3 interacts with MPK6 in a BiFC assay. BiFC was performed as described in Figure 1C. Coexpression of MKK3-nYFP and cYFP (Empty) served as a negative control (b), and coexpression of ZIP63 and ZIP63 (c) and CNX6 and CNX6 (d) served as positive controls. (**C**) The kinase-dead version of MPK6 (MPK6-kd) is phosphorylated by MKKK20-MKK3. Phosphorylation (top) and CBB-stained protein (bottom) are shown. (**D**) Presence of CRK21 further increases MPK6 phosphorylation. Phosphorylation (top) and CBB-stained protein (bottom) are shown. Kinase assays were performed as described in Figure 2. Images represent 1 out of 3 independent experiments. Scale bars, 50 μm.

### The MKKK20 signaling pathway is modulated by calmodulins and the protein phosphatase PP2C76

As shown in Figures 1B and 1C, numerous calmodulin and calmodulin-like proteins strongly interact with MKKK20 under stringent conditions. CaM7 showed the strongest interaction with MKKK20. *In vitro* kinase assays show that CaM7 was slightly phosphorylated by MKKK20 (Figure 5A; lane 6 and lane 8). In contrast, CRK21-CD and MKKK20 phosphorylation status showed no significant change with or without CaM7 (compare Figure 5A; lane 8 to lane 3 or 7). To further elucidate the effect of CaM7 on the CRK21 pathway, CaM7 was incubated with CRK21-CD, MKK20, and MKK3 in an *in-vitro* kinase assay (Figure 5B). When all four proteins were present in the same reaction, MKKK20 and MKK3 showed stronger phosphorylation (Figure 5B; lane 5) than in control assays (Figure 5B; lanes1-4). In addition, the phosphorylation of CaM7 was stronger in the presence of all three other proteins (Figure 5B; lane 5) as compared to MKKK20 and MKK3 alone (Figure 5B; lane 4), and CRK21 enhanced phosphorylation of MKKK20. This shows that CaM7 positively affects phosphorylation of MKKK20 and thus MKK3.

Phosphatases have been widely reported as negative modulators of MAPK signaling pathways (Anderson et al., 2011; Ghorbel et al., 2019; Lumbreras et al., 2010). PP2C76 (At5G53140) belongs to the PP2C family (clade F) of protein phosphatases. It was identified by a Y2H library screening for MKKK20 interactors (Benhamman *et al*., 2017). To explore the function of PP2C76 in the CRK21-MKKK20-MKK3-MPK6 cascade, we tested the interaction between PP2C76 and each member of the cascade as well as CaM7. Only MKKK20 was found to interact with PP2C76 in the Y2H assay (Figure 5C). In an *in vitro* kinase assay, MKKK20 autophosphorylation decreased with increasing amounts of PP2C76, indicating that PP2C76 dephosphorylated the autophosphorylated MKKK20 (Figure 5D and Figure S1; lane 3-5). Surprisingly, PP2C76 was also phosphorylated in the presence of MKKK20 (Figure 5D), but no PP2C76 phosphorylation was observed in the absence of MKKK20 (Figure 5D; lane 1 and Figure S1; lane 1). To show that the reduced autophosphorylation of MKKK20 is not due to the high concentration of protein present in the *in-vitro* assay, we co-incubated MKKK20 with 2 µg of BSA (Figure 5D; lanes 6 and 7). MKKK20 autophosphorylation was not reduced, indicating that MKKK20 was dephosphorylated by PP2C76. These results indicate that MKKK20 phosphorylates PP2C76, which dephosphorylates MKKK20 in turn, at least *in vitro*. It remains to be determined whether PP2C76 phosphorylation is required for its activity.

### Subcellular localization of proteins in the CRK21-initiated signaling pathway

To determine the localization of each protein in the CRK21 pathway, we transiently overexpressed the signaling proteins as fusions with the green fluorescent protein (GFP) in onion epidermal cells. As shown in Figure 6, the positive control, GFP alone, MKKK20-GFP, MKK3-GFP (Benhamman *et al*., 2017), MPK6-GFP, and CaM7-GFP were all prominently localized in the nucleus (Figure 6A, E, G, I, K). Some MAPKKKs are known to localize to the plasma membrane (Hashimoto-Sugimoto et al., 2016; Ozoe et al., 2002; Yamamoto et al., 2010), and MKKK20 also shows fluorescence at the plasma membrane. In contrast, PP2C76-GFP was mainly localized in the nucleus and in some discrete locales (possibly organelles) (Figure 6M), which is consistent with protein localization prediction tools (SUBA4: http://suba.live). CRK21, as a receptor kinase, is prominently located on the cell surface (Figure 6C).

**Figure 6.**
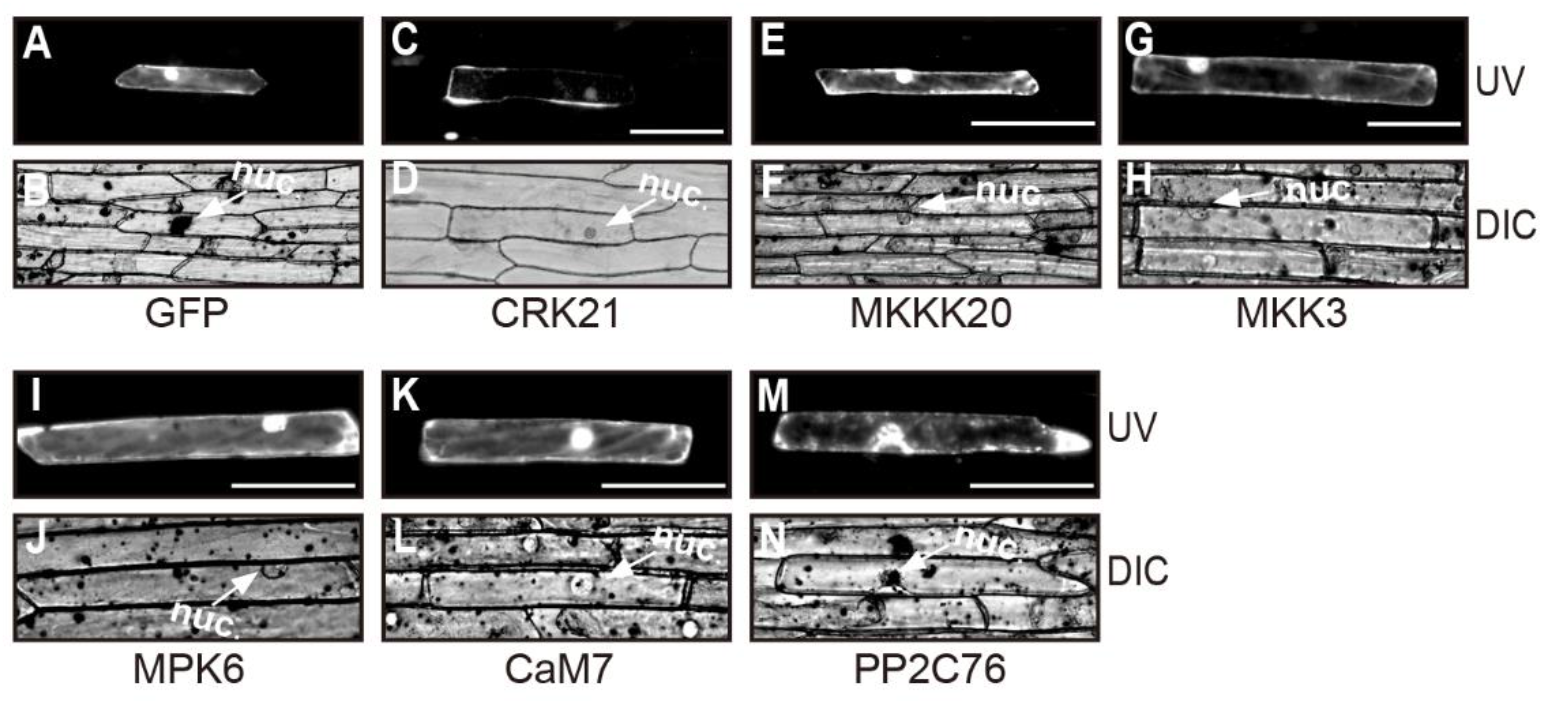
Protein subcellular localization in onion epidermal cells. Visualization of onion epidermal cells expressing GFP (**A, B**), CRK21-GFP (**C, D**), MKKK20-GFP (**E, F**), MKK3-GFP (**G, H**), MPK6-GFP (**I, J**), CaM7-GFP (**K, L**), and PP2C76-GFP (**M, N**). Images represent 1 out of 3 independent experiments. Scale bars, 100 μm.

### All members of the CRK21-MKKK20-MKK3-MPK6 signaling pathway and its CaM7 and PP2C76 modulators function in plant immunity

The CRKs CRK4, 5, 6, 10, 11, 19, 20, MKK3, and MPK6 function in plant immunity (Chen et al., 2004b; Doczi et al., 2007) (Asai *et al*., 2002) while MKKK20 is only known to function in responses to abiotic stress (see Introduction). To decipher the function of the CRK21-MKKK20-MKK3-MPK6 protein kinase cascade, T-DNA insertional mutant plants (*crk21-1*, *crk21-2*, *mkkk20-1*, *mkkk20-2*, *mkk3-1*, *mkk3-3, and mpk6-1*) were inoculated with *B. cinerea* and *P. syringae* pv. *tomato* (*Pst)* DC3000. All protein kinase mutant plants displayed reduced resistance to *B. cinerea* and *Pst* DC3000 compared with WT plants (Figure 7A-D) suggesting that each kinase in the CRK21-MKKK20-MKK3 cascade plays a role in immunity. The resistance of *mpk6* plants to pathogens was prominently weaker than resistance of the *crk21-1/2*, *mkkk20-1/2* and *mkk3*-*1/2* mutant plants (Figure 7A-D).

**Figure 7.**
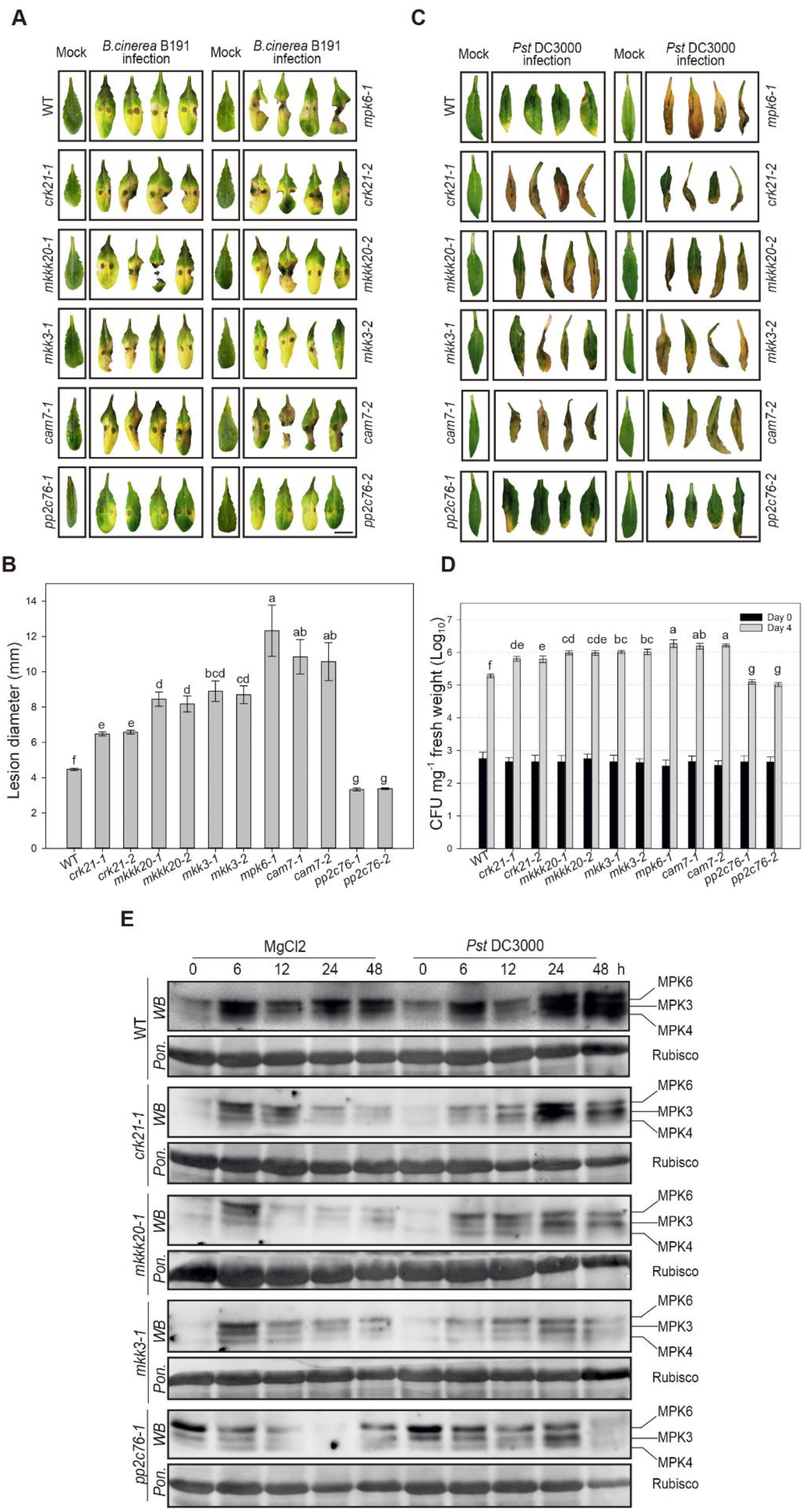
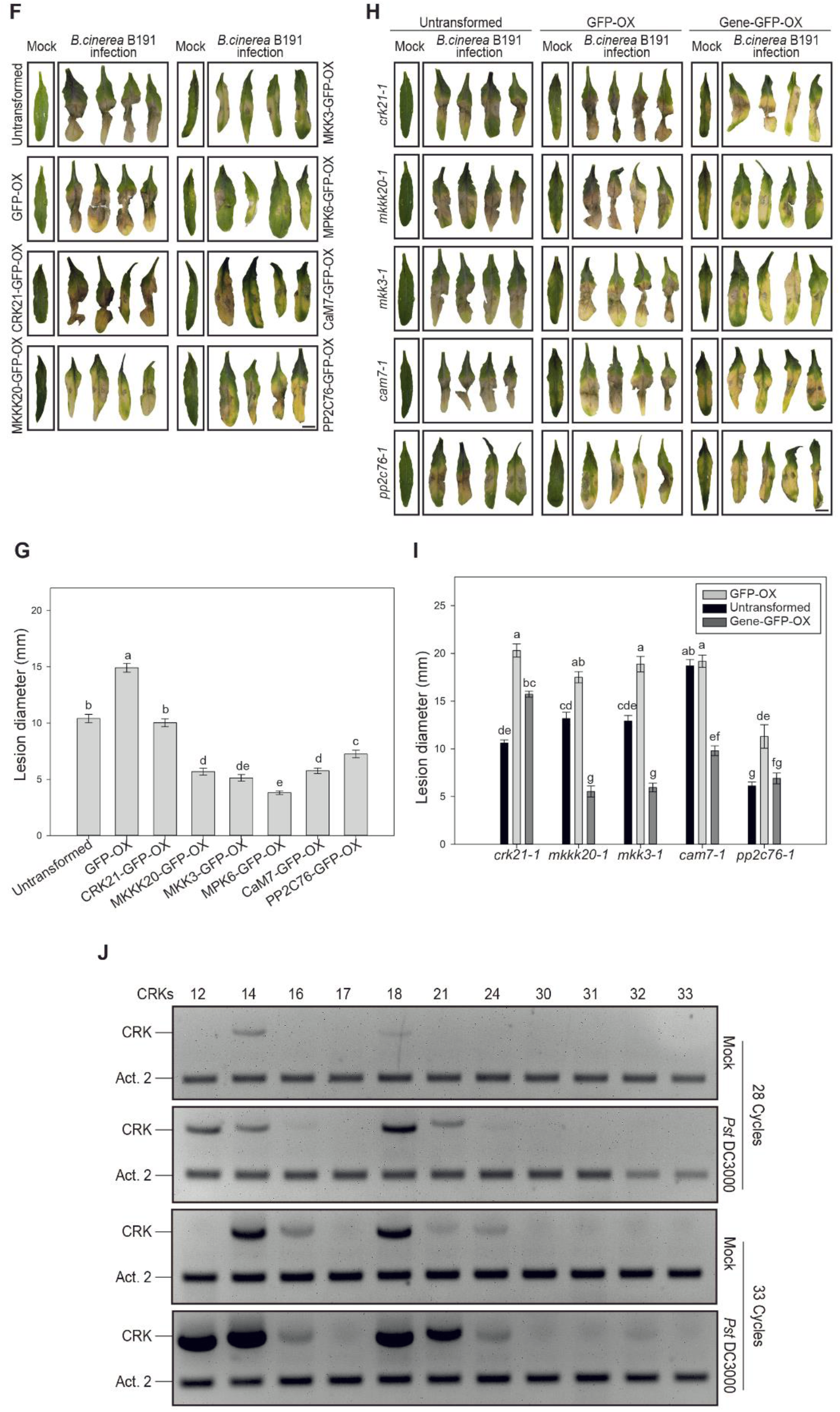
The CRKs-MKKK20-MKK3-MPK6 signaling pathway, CaM7, and PP2C76 function in plant immunity. (**A-D**) leaves from *crk21-1/2, mkkk20-1/2, mkk3-1/2, cam7-1/2, and mpk6* mutant plants exhibit enhanced disease symptoms, while *PP2C76-1/2* mutant plants exhibit reduced disease symptoms after *Botrytis cinerea* strain B191 (*B. cinerea* B191) infection (**A, B**) and *Pseudomonas syringae* pv. *tomato* DC3000 (EV) (*Pst* DC3000) (**C, D**) infection. (A) and (**C**) Disease symptoms of the leaves after 4 days (**A**) and 5 days (**C**) of infection. Images in (**A**) and (**C**) represent 4 representative leaves with varying degrees of symptoms out of 100-120 leaves (**A**) or 50-60 leaves (**C**) analyzed. (B) Statistical analysis of lesion diameters 4 days after inoculation with *B. cinerea* B191. (**D**) Statistical analysis of cfu of *Pst* DC3000 in leaves 4 days after infiltration. (**B**) and (**D**), Mean values ± s.d. of 100-120 (**B**) or 50-60 (**D**) leaves each from four independent experiments are shown. Different letters indicate statistically significant differences among treatments (*P* < 0.05). (**E**) Phosphorylation of MPK6 is decreased and/or delayed in leaves from *crk21-1*, *mkkk20-1*, and *mkk3-1* mutant plants after *Pst* DC3000 infiltration while in *pp2c76-1* mutant leaves, MPK6 is constitutively phosphorylated before infiltration. Phosphorylation of MPK3, MPK4, and MPK6 are shown. Images represent 1 out of 3 independent experiments. (**F-I**) Transient overexpression of the genes in the CRK21-MKKK20-MKK3-MPK6 pathway, including *CaM7* and *PP2C76*, increased the resistance to *B. cinerea* B191 infection. The full-length coding sequences of the corresponding genes were fused to the N-terminus of GFP (Gene-GFP) and were transiently overexpressed in 4-week-old WT (**F**) and mutant (**H**) Arabidopsis leaves through agroinfiltration (Gene-GFP-OX). The mutant leaves without any pretreatment (Untransformed) and transiently overexpressing GFP (GFP-OX) were used as negative controls. The leaves were detached 2 days after infiltration for *Botrytis cinerea* strain B191 infection. (**F**) and (**H**) Disease symptoms of WT (**F**) and mutant (**H**) leaves 3 days after infection with *B. cinerea* B191. Images represent 4 out of 30 leaves with varying degrees of symptoms. (**G**) and (**I**) Quantification and statistical analysis of lesion diameters in WT (**G**) and mutant (**I**) leaves 3 days after infection with *B. cinerea* B191. Mean values ± s.d. of five independent experiments measuring 30 leaves each are shown. Different letters indicate statistically significant differences among treatments in WT (**G**) or corresponding mutant leaves (**I**) (*P* < 0.05). Scale bars, 1 cm. (**J**) Semiquantitative RT-PCR analysis of CRK gene expression in leaves 2 days after infiltration with 10 mM MgCl_2_ (mock) or *Pst* DC3000. Images represent 1 out of 3 experiments.

To further confirm that the CRK21-MKKK20-MKK3 cascade is involved in resistance to pathogens, the phosphorylation status of MPK6 in WT and in *crk21-1*, *mkkk20-1*, and *mkk3-1* mutant leaves was monitored by western blot (WB) at different time points after *Pst* DC3000 infiltration (10^6^ cfu/mL in 10 mM MgCl_2_) (Figure 7E and S3). As expected, MPK6 phosphorylation in WT leaves was increased after MgCl_2_ infiltration due to wounding by infiltration (Ichimura et al., 2000), and it increased further at 24 and 48 h after *Pst* DC3000 infiltration (wounding and pathogen effects (Rasmussen et al., 2012)). Compared to WT plants, the *crk21-1*, *mkkk20-1*, and *mkk3-1* mutants showed generally weaker MAPK signals. These results suggest that upon *Pst* DC3000 infection, the activity of MPK6 is positively regulated by CRK21, MKKK20, and MKK3. The higher *Pst* DC3000-induced phosphorylation at 24 and 48 hpi was also observed in a former report (Asai *et al*., 2002). It is worth noting that the phosphorylation of the MAPKs was weaker in *crk21-1*, *mkkk20-1*, and *mkk3-1* MgCl_2_ injection controls compared to WT plants (Figure 7E and S3), suggesting that CRK21, MKKK20, and MKK3 also play key roles in the wound-activation of MPK6, MPK4, and MPK3.

In disease assays, the calmodulin mutants *cam7-1* and *cam7-2* were both sensitive to *Pst* DC3000 and *B. cinerea* infections, similar to *crk21-1/2*, *mkkk20-1/2*, and *mkk3-1* plants (Figure 7A-D). Since CaM7 increases phosphorylation of MKKK20 and MKK3, these results suggest that CaM7 plays a positive role in in plant immune responses via interaction with the MKKK20 signaling cascade.

In contrast, leaves from the phosphatase mutants *pp2c76-1* and *pp2c76-2* exhibited stronger resistance to both *B. cinerea* and *Pst* DC3000 than wild-type plants and other MKKK20-pathway mutants (Figure 7A-D). In contrast to WT and the other kinase mutant plants, *pp2c76-1* plants exhibited different MPK6 activation kinetics. MPK6 phosphorylation was high at time zero, regardless of treatment. After *Pst* DC3000 inoculation, MPK phosphorylation decreased slower than in the wounding controls (Figure 7E and S3). At later times, MAPK activity is first downregulated and then goes back to background level in wounded plants but decreases further at 48 hours after infection. Taken together, the results suggest that the CRK21-MKKK20-MKK3-MPK6 cascade contributes to fungal and bacterial pathogen resistance, and that PP2C76 is a negative modulator of this cascade.

### Complementation analysis of the CRK21-MKKK20-MKK3-MPK6 signaling pathway confirms its role in plant immunity

To further confirm the function of the *CRK21-MKKK20-MKK3-MPK6* signaling pathway in the plant immune response, we transiently overexpressed the *GFP*-tagged genes in leaves of Arabidopsis WT and of the corresponding mutants through agroinfiltration. *GFP*-overexpressing (GFP-OX) leaves were used as negative controls. Presence of recombinant proteins was shown through GFP fluorescence (Figure S3). *B. cinerea* infection in untransformed leaves resulted in a disease phenotype and cell death (Figure 7F, upper left panel), as compared to mock-inoculated leaves, and susceptibility to *B. cinerea* was further enhanced in leaves transformed with the *GFP* gene (Figure 7F and G). In contrast, when leaves were infiltrated with one of the six signaling genes fused to *GFP*, lesion diameters in *B. cinerea*-inoculated leaves were the same as in untransformed plants (*CRK21-GFP*) or smaller, and all were smaller as compared to *GFP*-overexpressing leaves. This indicates that all six genes positively regulate immunity to *B. cinerea* (Figure 7F and G). When the signaling genes as *GFP*-fusions or *GFP* alone were expressed in leaves of the corresponding mutant plants, the results were similar (Figure 7 H and I). All leaves overexpressing the five signaling genes (*MPK6* was not tested) in the corresponding mutants showed decreased lesion diameters as compared to mutant lines that expressed GFP alone, indicating that they positively regulate immunity. Figure 7H and 7I also confirm that *crk21-1*, *mkkk20-1*, *mkk3-1*, and *pp2c76-1* mutants are less susceptible to *B. cinerea* than *cam7-1* mutants (Figure 7A and B). Surprisingly, the overexpression of *PP2C76-GFP* did not increase lesion diameters, as expected. In Figure 7A and B, *pp2c76* mutants showed decreased lesion diameters (increased resistance) as compared to wildtype plants, consistent with a role of PP2C76 as a negative modulator of the pathway. Consistent with Figure 7A and B, the untransformed *pp2c76-1* mutant exhibited the smallest lesion diameters, but overexpression of *PP2C76-GFP* did not revert this effect. We did not include wild type plants in this experiment, but when compared to Figure 7A and B, the data indicate that *PP2C76-GFP* overexpression had no effect on resistance to *B. cinerea*. Perhaps, the GFP fusion reduced phosphatase activity of PP2C76. Taken together, these results indicate that the signaling pathway containing CRK21, MKKK20, MKK3, MPK6, PP2C76, and CaM7 is involved in defenses against *B. cinerea* B191.

### Several CRKs function in plant immune responses

Beside CRK21, several CRKs interact and phosphorylate MKKK20 (Figure 3). Our data show that *CRK21* participates in pathogen resistance, however *crk21-1/2* mutant plants are not as susceptible toward *Pst* DC3000 and *B. cinerea* as *mkkk20*, *mkk3*, or *mpk6* mutant plants (Figure 7 A-D), suggesting a possible redundancy of CRKs acting upstream of *MKKK20*.

To test if these MKKK20-interacting CRKs were also involved in plant immune responses, their gene expression was determined in mock- and *Pst* DC3000-infiltrated Arabidopsis leaves by semiquantitative RT-PCR. Of the 11 CRKs tested (Figure 7J), five CRKs that interact with MKKK20 (CRK14, 16, 18, 21 and 24) were expressed in mock-treated leaves, and their expression was increased in response to *Pst* DC3000 infection. CRK12 was only expressed in *Pst*-treated leaves, while CRKs 30, 31, 32, and 33 were neither expressed in mock nor *Pst* DC3000-infiltrated leaves, although CRK30 and CRK33 interacted with MKKK20 (Figure 3A and B). CRK expression in response to *Pst* DC3000 infection maybe a positive feedback mechanism.

## Discussion

We provide evidence for a novel MAPK cascade, consisting of MKKK20, MKK3, and MPK6, that functions in the response to the hemibiotrophic bacterial pathogen *Pst* DC3000 and the necrotrophic fungal pathogen *B. cinerea*. In addition, we demonstrate *in vitro* and *in vivo* that a cysteine-rich receptor-like kinase (CRK21), a calmodulin (CaM7) and a PP2C type phosphatase (PP2C76) are MKKK20-interacting proteins that all function in plant immunity.

In a search for potential activators of MKKK20, we identified CRK21 and demonstrated that CRK21 activates the MKKK20-MKK3-MPK6 cascade. It interacted with MKKK20 in Y2H and BiFC assays (Figure 1B and Figure 1C) and phosphorylated MKKK20, which consequently increased the phosphorylation of downstream MKK3 and MPK6 *in vitro* (Figure 2A and B, Figure 4C and D). Based on our knowledge, there is only one additional MKKK known that is activated directly by an RLK through phosphorylation. Under cold stress, CRLK1 interacts with and directly activates MEKK1 through phosphorylation (Furuya *et al*., 2013; Yang *et al*., 2010a; Yang *et al*., 2010b). A better-known mechanism for activation of MKKKs is via receptor-like cytoplasmic kinases (RLCKs), which can be activated by extracellular RLKs (Cui et al., 2018; Jose et al., 2020).

The null mutant plants *crk21-1/2* showed weaker resistance to *B. cinerea* and *Pst* DC3000 than WT (Figure 7A-D), and overexpression of CRK21 strengthened the resistance to *B. cinerea* in both *crk21-1* mutant and WT plants (Figure 7F-I). Moreover, the *Pst* DC3000-induced MPK6 phosphorylation kinetics were different in the *crk21-1/2* null mutant plants as compared to that in WT plants (Figure 7E). However, the loss of resistance to *B. cinerea* and *Pst* DC3000 in *crk21-1/2* plants as compared to WT plants is less pronounced than in *mkkk20-1/2* and *mkk3-1/2* plants (Figure 7A-D). Similarly, increased resistance to *B. cinerea* in the CRK21 over-expressing plants was less pronounced than in MKKK20- and MKK3-expressing plants (Figure 7F-I), indicating that the CRK21 function could be complemented by functionally similar proteins. In a similar scenario, only triple mutants of the flagellin-induced CRK22, 28 and 29 showed reduced resistance to *P. syringae* (Yadeta *et al*., 2017). Therefore, we hypothesized that MKKK20 may also receive signals from other CRKs and we found that six additional CRKs interacted with MKKK20 (Figure 3A and B). While we were unable to show that these CRKs phosphorylate MKKK20 due to MKKK20 autophosphorylation, we found that MKKK20 phosphorylates five of the interacting CRKs (Figure 3D). The data are consistent with a reciprocal phosphorylation between CRKs and MKKK20. To understand the signaling dynamics, it will be important to define the series of phosphorylation events. In contrast to the other CRKs, CRK30 strongly inhibited the autophosphorylation of MKKK20 in kinase assays (Figure 3D; lane 7) indicating that CRK30 may be a negative regulator of MKKK20-mediated signaling pathways. Gene expression analysis indicated that CRK12, CRK14, CRK18, and CRK21 strongly respond to *Pst* DC3000 infection (Figure 7J) indicating that these CRKs also play a role in immunity and probably complement each other to achieve a robust immune response through the MKKK20 pathway. These results also indicate a positive feedback loop that promotes the recruitment of CRKs, presumably to strengthen the immune response. CRK30 was not induced by *Pst* DC3000, indicating that competing signals that activate CRK30 downregulate immune responses.

C group MAPKs (MPK1/2/7/14) as well as an A group (MPK6) and a D group (MPK8) MAPK act downstream of MKK3 to regulate a range of responses (Colcombet and Hirt, 2008; Jagodzik et al., 2018; Kumar *et al*., 2020). The cascade consisting of MKKK17/18, MKK3, and MPK7 modulates responses to ABA (Danquah et al., 2015), and the MKKK14-MKK3-MPK2 cascade plays a role in the wound-induced JA response. In addition, all members of the clade-III MAP3Ks (MKKK13-20) phosphorylate MPK2 through MKK3 in mesophyll protoplasts (Sozen *et al*., 2020). Our results establish a novel role for MKK3 as part of the MKKK20-MKK3-MPK6 cascade and confirm that MKK3 is an important convergence and divergence node in the plant stress signaling network.

The protein phosphatase 2C (PP2C) family has 80 members (Fuchs et al., 2013). The subclades A, B, C, D, F and G contain MAPK phosphatases that function in plant defenses (Carrasco et al., 2014; Couto et al., 2016; Lim et al., 2014; Ludwikow et al., 2014; Mine *et al*., 2017; Shubchynskyy *et al*., 2017; Sidonskaya et al., 2016). In clade F, PIA1 (At2g20630) and WIN2 (At4g31750) function in the response to *P. syringa*e (Lee et al., 2008b; Widjaja et al., 2010). We found that another member of the F-clade, PP2C76 (At5G53140), interacts with MKKK20 in Y2H (Figure 5C) and BiFC assays (Figure 1C; i), and that it negatively regulates the activity of MKKK20 and MPK3/6 (Figure 5D and Figure 7E). PP2C76 is in a subclade of clade F different from the subclade that contains PIA1 and WIN2 (Schweighofer *et al*., 2004). The *pp2c76-1/2* null mutant plants showed the strongest resistance to *B. cinerea* and *Pst3000* infection as compared to the other mutants we tested (Figure 7A-D). Our findings suggests that PP2C76 plays an important role in controlling the activity of MKKK20. Its activity must be tightly regulated by pathogen-related signals to allow for activation of MKKK20 during pathogen infection and to prevent MKKK20 from autoactivation in the absence of a stimulus. To our knowledge, only one other protein phosphatase that targets a MAPKKK has been reported in plants.

The Arabidopsis PP2C ABSCISIC ACID INSENSITIVE 1 (ABI1) negatively regulates abscisic acid signaling via dephosphorylation of MKKK20, which was named ABA-INSENSITIVE PROTEIN KINASE 1 (AIK1), and ABI1 restricted MKKK18 and MKKK20 activity *in vitro* (Li *et al*., 2017; Mitula et al., 2015).

While PP2C76 is a negative modulator of the CRK21-MKKK20-MKK3-MPK6 cascade, we identified MKKK20-interacting calmodulins as positive modulators of this pathway. Rapid increases in cytosolic Ca^2+^concentrations are universal signaling processes involved in a wide range of stress responses and cellular functions. Changes in cytosolic calcium concentrations can be input-specific and the ‘calcium signature’ can be decoded by Ca^2+^-binding proteins that function as Ca^2+^ sensors and transmitters by interacting with other proteins and modulating their activity (Kudla et al., 2010). They may also assume a bridge function that brings two proteins into proximity (Villalobo *et al*., 2017). Surprisingly, five of the seven Arabidopsis CaMs, as well as one CML, interacted with MKKK20 in Y2H and BiFC assays, with CaM7 showing the strongest interaction (Figure 1B and Figure 1C). CaM7 is a multifaceted protein functioning in a wide range of cellular processes such as responses to stress, pathogens, and photomorphogenesis. It can interact with proteins as diverse as the ABC transporter PEN3, cyclic nucleotide-gated channels, the MAPK MPK8, and the E3 ubiquitin ligase COP1 and directly regulate gene expression (Basu et al., 2021). Arabidopsis *cam7* and *cam7/pen3* mutants are susceptible to the non-adapted fungal pathogens *Phakopsora pachyrhizi* and *Blumeria graminis* f.sp. *hordei* (Campe et al., 2016). Here, we show that CaM7 enhances the activity of the CRK21-MKKK20-MKK3 cascade *in vitro* (Figure 5B), and that loss of CaM7 in *cam7-1/2* mutant plants resulted in high susceptibility to *B. cinerea* and *Pst* DC3000 (Figure 7A-D). This suggests that CaM7 plays an important positive role in the plant immune response.

As mentioned above, we showed that the CRK21-MKKK20-MKK3-MPK6 cascade is involved in responses to the model pathogens *Pst* DC3000 and *B. cinerea*. Phenotypical analysis of the infected mutant plants *crk21*, *mkkk20*, *mkk3*, *cam7*, as well as *pp2c76* confirmed our hypothesis. Compared to *mpk6* mutant plants, *crk21-1/2*, *mkkk20-1/2*, and *mkk3-1/2* were less susceptible to *B. cinerea* and *Pst* DC3000 infection (Figure 7A-D), consistent with reports showing that *MPK6* is involved in other MAPK cascades such as the well-known FLS2-MEKK1-MKK4/5-MPK3/6 cascade (Asai *et al*., 2002). This is in line with our MAPK phosphorylation assays showing that MPK6 remained active after pathogen infection in *crk21-1*, *mkkk20-1*, and *mkk3-1* mutant plants, albeit weaker than in WT plants (Figure 7E). The role of the MKKK20 cascade in immunity was further confirmed in disease assays with plants overexpressing the respective genes in both WT and mutant plants (Figure 7F-I). In these assays, agro-infiltration with the GFP construct alone (GFP-OX) significantly decreased the resistance of the plants, possibly because *Agrobacterium* is a plant pathogen itself (Figure 7F-I). Therefore, all lines that overexpress our genes of interest were compared to the GFP-OX line. Surprisingly, the over-expression of PP2C76 did not impair but significantly promoted the resistance to *B. cinerea* infection (Figure 7 F-I), which indicates that PP2C76 also functions as a positive modulator of other proteins that function in immunity.

As summarized in the model shown in Figure S5, this work uncovered a novel MAPK cascade, consisting of MKKK20, MKK3, and MPK6, that functions in resistance to diverse pathogens in Arabidopsis. The cascade is most likely activated in response to pathogen attack by the receptor kinase CRK21, and it is modulated positively by CaM7 and negatively by PP2C76. Additional CRKs (CRK12, 14, 16, 18, 24) also interact with MKKK20 and are induced by pathogen attack (Figs 3 and 7j). Unlike these CRKs, CRK30 may function as a negative regulator of MKKK20. A critical gap in this model is the lack of a pathogen-associated signal that activates the CRK receptor kinases. The CRKs identified here could also play a role in MAMP-triggered immunity through interaction with PRRs. The CRKs CRK4, CRK6, and CRK36 are known to interact with FLS2.

## Materials and Methods

### Plants materials and pathogens

The *A. thaliana* seeds of wild type (Col-0) and the homozygous T-DNA lines (Table S2) were ordered from the Arabidopsis Biological Resource Center (ABRC) and grown in soil under a long day light regime at 23°C. T-DNA insertional mutants were identified based on information on TAIR (https://www.arabidopsis.org). Homozygosity was confirmed by PCR with the primers listed in Table S2. The *mkkk20-1* (Kim *et al*., 2012), *mkkk20-2* (Li *et al*., 2017), *mkk3-1* (Lee, 2015), *cam7-2* (Campe *et al*., 2016) and *mpk6-1* (Liu and Zhang, 2004) mutants were characterized as described in the above references. The expression of *CRK21*, *MKK3*, *CaM7*, and *PP2C76* in WT (Col-0) and the corresponding mutants were verified by RT-PCR with the primers listed in Table S3 (Figure S4). The *mkk3-2* mutant showed residual *MKK3* transcript (Figure S4B). For RT-PCR, total RNA was extracted from 4-week-old soil-grown plants by TRIzol (Invitrogen), reverse transcribed by the M-MLV RTase cDNA synthesis kit from Invitrogen, and cDNA was amplified for 30 (Figure S4B-D) and 35 (Figure S4A) cycles.

The pathogens *Pseudomonas syringae* pv. *tomato DC3000* (*Pst* DC3000) and *Botrytis cinerea* strain B191 were gifts from professor Kamal Bouarab (Biology department, Université de Sherbrooke, Canada).

### cDNA library preparation and constructs

Total RNA from whole plant was isolated using TRIzol (Invitrogen) from either 7-day-old Col-0 (WT) seedlings grown on plates containing half-strength Murashige and Skoog (MS) and 1% sucrose, or from 4-week-old leaves that were treated with *Pst* DC3000 3 days before harvesting. The M-MLV RTase cDNA synthesis kit from Invitrogen was used for reverse transcription.

The *A. thaliana* cDNAs encoding the full ORF of MKKK20 (AT3G50310), MKK3 (AT5G40440), MPK6 (AT2G43790), PP2C76 (AT5G53140), CML10 (AT2G41090), CaM1 (AT5G37780), CaM4 (AT1G66410), CaM9 (AT3G51920), CaM7 (AT3G43810), CaM6 (AT5G21274), Ca^2+^ Binding EF-hand protein (AT5G28900) were all obtained by PCR from the *A. thaliana* whole plant cDNA library. The cDNAs encoding the full ORF (for localization analysis and BiFC) and the cytosolic domains (for protein expression and Y2H) of CRK12 (AT4G23200), CRK14 (AT4G23220), CRK16 (AT4G23240), CRK17 (AT4G23250), CRK18 (AT4G23260), CRK21 (AT4G23290), CRK24 (AT4G23320), CRK30 (AT4G11460), CRK32 (AT4G11480), CRK33 (AT4G11490) were acquired by PCR from the *Pst* DC3000 treated leaf cDNA library (primers for the cytosolic domains are shown in Table S1). The PCR products were cloned into pQLinkHD and pDEST565 to express and purify 6xHis- and GST-Tagged proteins; into pDEST22 and pDEST32 (Invitrogen) for Y2H; into pMDC83 for subcellular localization analysis; into pSPYNE-35S^GW^ and pSPYCE-35S^GW^ for BiFC through gateway cloning system (Invitrogen). The resulting plasmids were verified by sequencing.

### Yeast two-hybrid assay (Y2H)

Y2H assays were performed as described (Benhamman *et al*., 2017) using the Gateway^TM^ yeast two-hybrid bait (pDEST32) and prey (pDEST22) vectors from Invitrogen^TM^. In addition to the interaction controls from the kits (C^s^: pEXPTM32-Krev1/pEXPTM22-RalGDS-wt as a strong positive interaction control; C^w^: pEXPTM32-Krev1/pEXPTM22-RalGDS-m1 as a weak positive interaction control; C^0a^: pEXPTM32-Krev1/pEXPTM22-RalGDS-m2 as a negative interaction control), pDEST32-Empty/pDEST22-Empty (C^0b^) and pDEST32-bait/pDEST22-Empty were also used as the second and third negative interaction controls.

### Bimolecular fluorescence complementation assay (BiFC)

The full-length coding sequence of the genes were cloned into pSPYNE-35S^GW^ (N-terminal YFP) and/or pSPYCE-35S^GW^ (C-terminal YFP) through gateway cloning (Invitrogen™). BiFC was carried out as described (Gou et al., 2015) using transformed *A. tumefaciens* GV3101 for infiltration into wild type *N. benthamiana* leaves. The fluorescence of the transformed epidermal cells was observed 2-3 days after infiltration.

### Subcellular localization analysis

The constructs pMDC83 containing the genes of interest were transiently transformed into spring onion epidermal cells by microparticle bombardment using a Bio-Rad PDS-1000/He bombardment system as described (Benhamman *et al*., 2017).

### *In vitro* kinase assay

Protein expression and purification was performed as described (Benhamman *et al*., 2017). About 1 µg of each 6xHis- or GST-tagged protein was put in 35 µl kinase reaction buffer (50 mM Tris–HCl, pH 7.5, 1 mM DTT, 10 mM MgCl_2_, 1.5 mM MnCl_2_, 50 µM ATP, 2 mM CaCl_2_, and 0.037 MBq [γ-^32^P] ATP). After 30 min incubation at 30°C, the reactions were stopped by adding 5 µl of 6X SDS sample loading buffer and heating at 95°C for 5 min.

After SDS-PAGE, protein bands and radioactive bands were detected with an ImageQuant LA4000 imaging system (GE Healthcare) and a Typhoon Trio phosphorimager (GE Healthcare), respectively.

### Pathogen treatment

Treatment of Arabidopsis with *Pst* DC3000 (pressure infiltration) and *B. cinerea* B191 (drop inoculation) were performed as described (Ingle and Roden, 2014). The *Pst* DC3000 infected leaves were harvested and weighted 4 days post-infection (dpi). The leaves were ground in 10 mM MgCl_2_ with the final concentration of 1 mg/ml after surface sterilization by immersing in 70% ethanol for 30 s and rinsing in distilled water. The resulting suspensions were diluted by 10 mM MgCl_2_ to generate a serial dilution (from 10^-2^ to 10^-6^) and subsequently plated onto LB plate containing 20 µg/ml rifampicin and 50 µg/ml kanamycin. The colonies were counted 2 days after incubation in the dark at 28°C. 3-4 days after inoculation of the detached Arabidopsis leaves with *B. cinerea* (23°C with 12 h photoperiod), the lesion diameters were measured as the length of the diagonal, which goes from an upper corner across the leaf to the lower corner of the necrotic tissue.

### Complementation Test

The genes of interest were cloned into the plasmid pMDC83, and transiently overexpressed in 4-week-old WT and mutant leaves through agroinfiltration as described above (see BiFC).

### Protein extraction and immunoblot analysis

The MgCl_2_ or *Pst* DC3000 treated leaves from WT and *crk21-1*, *mkkk20-1*, *mkk3-1*, *cam7-1*, and *PP2C76-1* mutant plants were detached at indicated hours post-inoculation (hpi) with *Pst* DC3000 in 10 mM MgCl_2_ or with 10 mM MgCl_2_ alone (control) and ground in liquid nitrogen. Total protein extraction was carried out as described (Ichimura *et al*., 2000). Thirty (30) µg total proteins of each sample were separated by 10% SDS-PAGE, subsequently blotted onto a nitrocellulose membrane (Millipore 0.2 µM), and proteins visualized by staining with 0.1 % ponceau S. The rubisco band was used to verify equal protein loading and transfer. To render the MAPK kinetics comparable among the different lines, proteins in all samples were separated by SDS-PAGE at the same time and under the same conditions, but on separate gels. Proteins from all the gels were then transferred to one membrane, as shown in Figure S2. The primary antibody was anti-p-ERK (anti-pERK MAPK [Phospho-p44/p42 MAPK, ERK1/2, Thr202/Tyr204, D13.14.4E], Cell Signaling Technology) that specifically recognizes the [T-X-Y] MAPK activation motif. The secondary antibody was HRP conjugated Goat anti-Rabbit IgG(H+L) (Thermo Fisher Scientific).

### Semi-quantitative RT-PCR

Leaves were harvested 24 h after treatment and total RNA was purified as described above. RT-PCR was performed with 10 ng of total RNA using OneTaq (New England Biolabs) PCR reaction system with the primers listed in Table S3. PCR was run for 28 and 33 cycles. Constitutively expressed *ACTIN2* (AT3G18780) was co-amplified with each sample and used as an internal standard. RT-PCR products were visualized using a 1.3 % agarose gel.

### Funding

This work was supported by the Natural Sciences and Engineering Research Council of Canada RGPIN-2019-05931 (to D.P.M.); The China Scholarship Council (CSC) (to F.B.).

### Author contributions

Conceptualization, F.B. and D.P.M.; Methodology, F.B. and D.P.M.; Investigation, F.B.; Writing – Original Draft, F.B. and J.W.S.; Writing – Review & Editing, F.B., J.W.S., and D.P.M.; Funding Acquisition, F.B. and D.P.M.; Resources, D.P.M.; Supervision, F.B. and D.P.M.

## Acknowledgments

We thank professor Kamal Bouarab for kindly providing *Pseudomonas syringae* pv. *tomato DC3000* (*Pst* DC3000) and *Botrytis cinerea* strain B191. No conflict of interest declared.

