## Supplemental Figures S1 to S5 Tables S1 to S3 for "A complete MAPK cascade, a calmodulin, and a protein phosphatase act downstream of CRK receptor kinases and regulate *Arabidopsis* innate immunity"

**The supplemental materials include:**

Figures S1 to S5

Tables S1 to S3

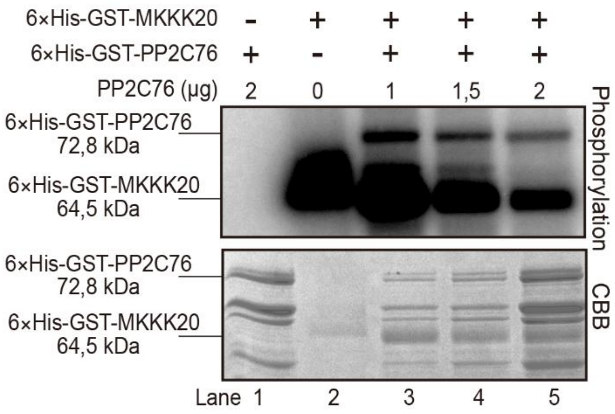


**Figure S1. PP2C76 dephosphorylates MKKK20 in an *in vitro* kinase assay.**
Phosphorylation (top) and proteins (bottom) are shown. 6xHis-GST-tagged proteins were expressed in bacteria and purified. Proteins were combined as shown and incubated in kinase assay buffer containing [$\gamma$-$^{32}$P]-ATP. CBB, Coomassie Brilliant Blue stain.

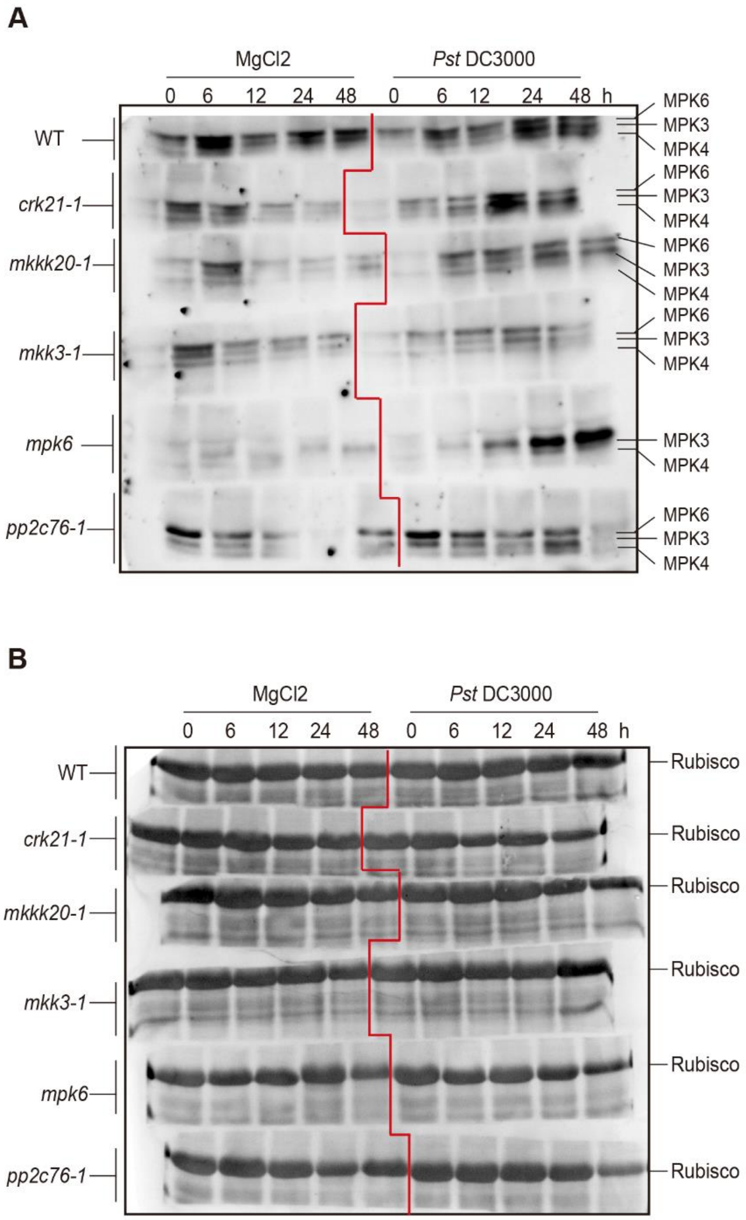


**Figure S2. Phosphorylation of MPK6 is regulated by CRK21-MKKK20-MKK3 and PP2C76.** (**A**) Total proteins were extracted from the wild-type (WT) and mutant leaves and collected at indicated hours post-inoculation (hpi) with either *Pst* DC3000 in 10 mM $MgCl_2$ or 10 mM $MgCl_2$ (control). An immunoblot is shown with an antibody against the phosphorylated TEY activation motif of MPK3, MPK4, and MPK6. After SDS-PAGE, all samples shown were transferred onto one nitrocellulose membrane. (**B**) Ponceau S staining of the membrane was used to verify equal loading of total protein. The position of MPK3, MPK4, MPK6 was based on the *mpk6* samples and their MW. To render the MAPK kinetics comparable among the different lines, proteins in all samples were separated by SDS-PAGE at the same time and under the same conditions, but on separate gels. Proteins from all the gels were then transferred to one membrane. For Figure 7E, the different mutant lines were aligned with the WT line.

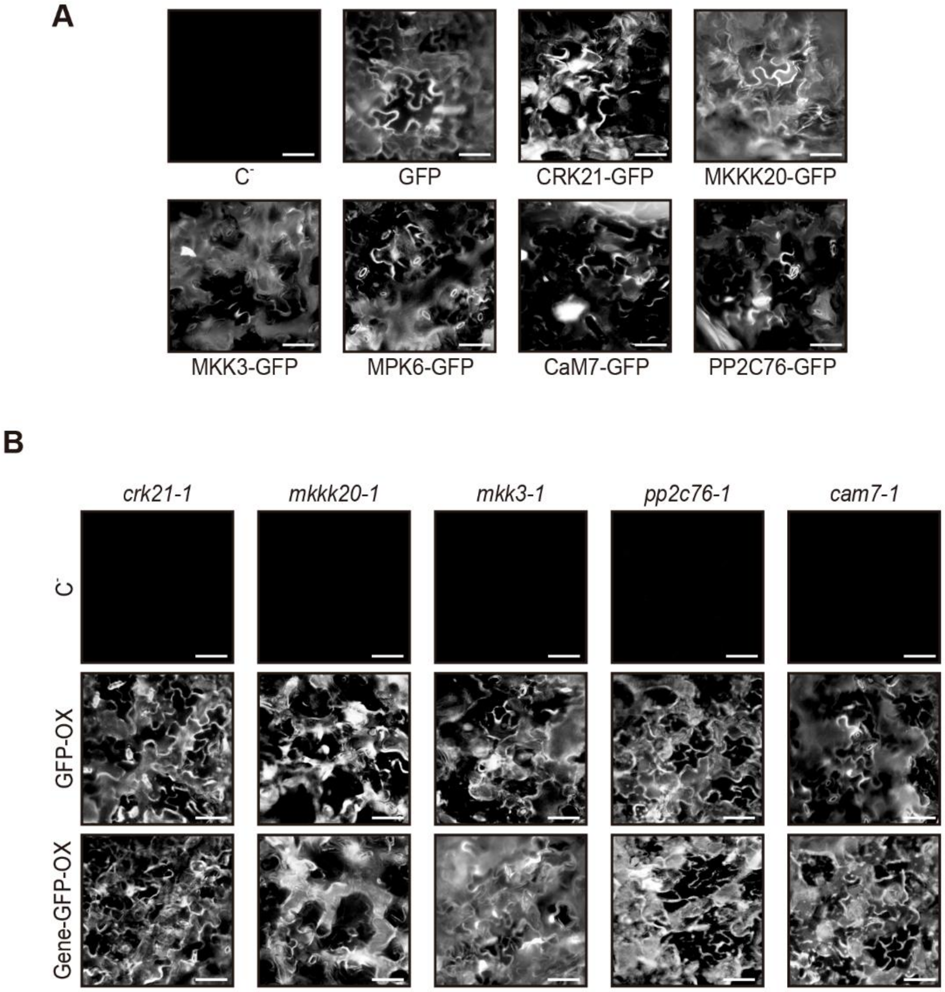


**Figure S3. Expression verification of GFP and GFP-tagged proteins in Arabidopsis.** The proteins GFP, CRK21-GFP, MKKK20-GFP, MKK3-GFP, MPK6-GFP, PP2C76-GFP, and CaM7-GFP were transiently overexpressed in WT (**A**) and the corresponding mutant plants (**B**) (Gene-GFP-OX) through agroinfiltration. 2 days after infiltration, the leaves were detached and observed under a fluorescence microscope. Leaves were infiltrated with P19, the RNA silencing suppressor from tomato bushy stunt virus, and with GFP alone as negative controls (C$^-$; GFP-OX). Scale bars, 50 µm.

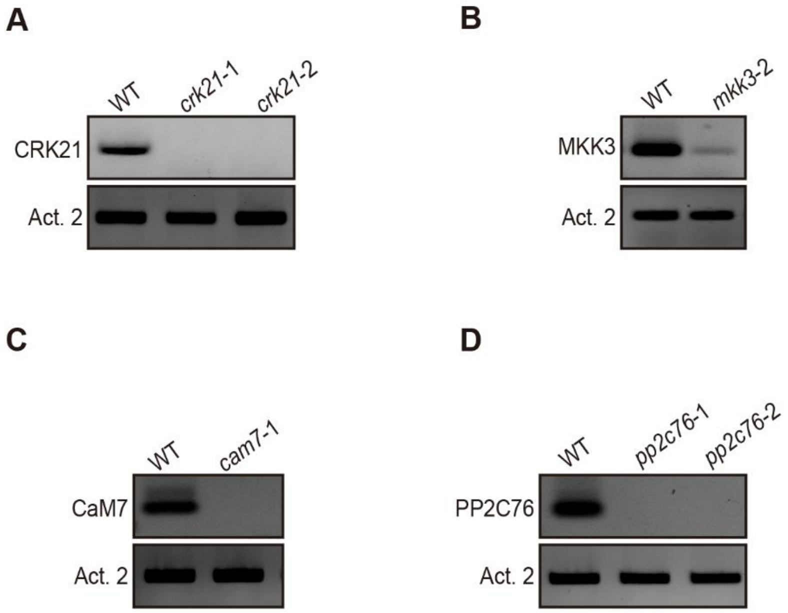


**Figure S4. Semiquantitative RT-PCR analysis of the gene expression in WT and T-DNA insertion lines.** The expression of *CRK21* (**A**), *MKK3* (**B**), *CaM7* (**C**), and *PP2C76* (**D**) (top) and *Actin2* (bottom) in WT (Col-0) and corresponding mutants are shown.

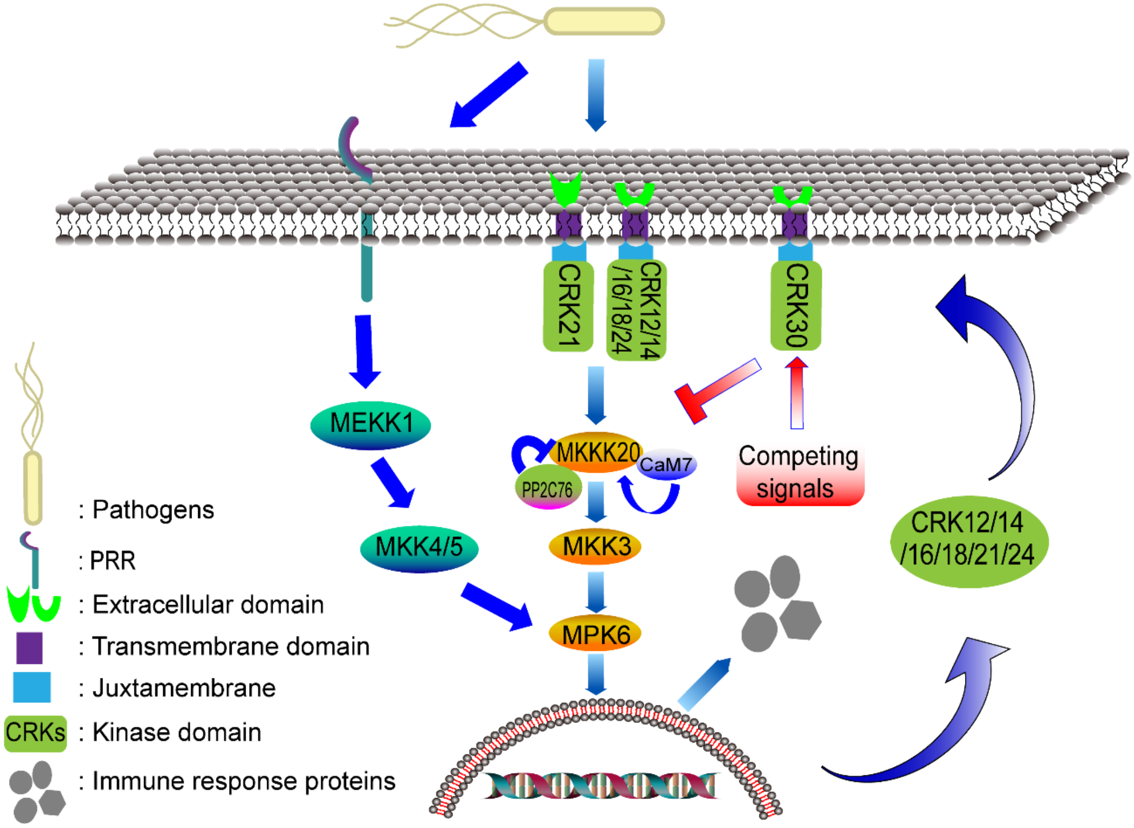


**Figure S5. Model of CRK induced signaling in Arabidopsis.** Unknown signals derived from pathogens or other stresses activate CRKs, which interact with other CRKs resulting in activation or inhibition of MAPK pathways. The activity of the MKKK20 cascade is modulated by the protein phosphatase PP2C76 and calmodulins such as CaM7. The result is immunity to fungal and bacterial pathogens as well as a positive feedback regulation through CRK expression. Other pathogen-derived signals such as MAMPs activate parallel signaling pathways that induce MAMP-triggered immunity via MEKK1 and MKK4/5. The MEKK1 and the MKKK20 pathways converge on MPK6 which activates defense gene expression. MAMP-triggered immunity and the immune response induced by the CRK-initiated pathway may be coordinated to activate a robust immune response.

**Table S1. Gateway primers for the cytosolic domains of CRKs.**

| CRKs | Locus ID | Primer (Forward) (5’ to 3’) | Primer (Reverse) (5’ to 3’) |
|---|---|---|---|
| CRK 12 | AT4G23200 | *GGGGACAAGTTTGTACAAAAAAGCAGGCTTC*GGTATTACGACTCTACACTTCCAACAAC | *GGGGACCACTTTGTACAAGAAAGCTGGGTC*TCAACGAGGATCTAGATTTGTAATGG |
| CRK 14 | AT4G23220 | *GGGGACAAGTTTGTACAAAAAAGCAGGCTTC*TTTGTTGTTTATAGGAGGAGGAAATCATAC | *GGGGACCACTTTGTACAAGAAAGCTGGGTC*TCAACGAGGTTCAAAATCTGTAATGGTTAC |
| CRK 16 | AT4G23240 | *GGGGACAAGTTTGTACAAAAAAGCAGGCTTC*AGGAGGAGAAAAGCCTACCAAGAGTTTG | *GGGGACCACTTTGTACAAGAAAGCTGGGTC*TCAACGAAGATCAACACTCGTGATCGATG |
| CRK 17 | AT4G23250 | *GGGGACAAGTTTGTACAAAAAAGCAGGCTTC*TGCAAGAGAAGAAAGCAAAAGCAAG | *GGGGACCACTTTGTACAAGAAAGCTGGGTC*TCAATTTCTTGTAGAAATCTTTCCGCCATC |
| CRK 18 | AT4G23260 | *GGGGACAAGTTTGTACAAAAAAGCAGGCTTC*AGAAGAAAGCAAAAGCAAGAAATGGATC | *GGGGACCACTTTGTACAAGAAAGCTGGGTC*TCAACGAGGATTAACGTCCGTGATTG |
| CRK 21 | AT4G23290 | *GGGGACAAGTTTGTACAAAAAAGCAGGCTTC*TCTAGGAGGAGAAAAGCTTACCAATC | *GGGGACCACTTTGTACAAGAAAGCTGGGTC*TTAACGAGGTCTAACACTCGTGATC |
| CRK 24 | AT4G23320 | *GGGGACAAGTTTGTACAAAAAAGCAGGCTTC*TGGAAGAGGAGAAAAGCATATAAAACAAAG | *GGGGACCACTTTGTACAAGAAAGCTGGGTC*TTAACGAGGACTAACACACGTGACCGATAC |
| CRK 30 | AT4G11460 | *GGGGACAAGTTTGTACAAAAAAGCAGGCTTC*TGCAGGAGTAGAAAAAAATATCAAGC | *GGGGACCACTTTGTACAAGAAAGCTGGGTC*TCAATCCTCAGTGTTTCTATACATTGAC |
| CRK 32 | AT4G11480 | *GGGGACAAGTTTGTACAAAAAAGCAGGCTTC*CGGAAGAGAAGACAATCGTACAAAAC | *GGGGACCACTTTGTACAAGAAAGCTGGGTC*TTACCGAGGAGTGACTCTAGTGATTG |
| CRK 33 | AT4G11490 | *GGGGACAAGTTTGTACAAAAAAGCAGGCTTC*GTTTGCAGAAAGAGAAAAACTGATC | *GGGGACCACTTTGTACAAGAAAGCTGGGTC*TCAGCGAGGAACTAAGTCATCAATC |

*Note: Italic* nucleobases are attB sequence.

**Table S2. T-DNA insertion lines.**

| Lines | Locus | FST-ID/Stock ID | Ecotype | LP (Left genomic primer) (5' to 3') | RP (Right genomic primer) (5' to 3') |
|---|---|---|---|---|---|
| *crk21-1* | AT4G23290 | SALK_035259 | Columbia | CGCTGTAAGATCATCCGTAGC | TTGATTAATGGATCGCTCGAC |
| *crk21-2* | AT4G23290 | SALK_022512C | Columbia | AAAAGAAGGGAGGAATGAGTCC | GATATATTTTTGCCATCTGCGC |
| *mkkk20-1* | AT3G50310 | SALK_124389 | Columbia | ACACCACAAACCTCAACTTGC | AAGCTTCCTCTCGAAGCGTAC |
| *mkkk20-2* | AT3G50310 | SALK_124398 | Columbia | TAAACATGCAAATTGCAGCTG | TCAGAATTGGAATGGGAATTG |
| *mkk3-1* | AT5G40440 | SALK_051970 | Columbia | GTTTGTGGGGTTTTTCACATG | CTTCACCAACACCACCAAAAC |
| *mkk3-2* | AT5G40440 | CS872767/SAIL_78_G03 | Columbia | TTTGTTCCCCATGTCTACGTC | CGTTTGGATCCAAAGATGAAG |
| *mpk6-1* | AT2G43790 | CS31099 | Wassilewskija | GCGGATCCATGGACGGTGGTTCAAGG | GCACTAGTCTATTGCTGATATTCTGGATTG |
| *cam7-1* | AT3G43810 | CS834151 | Columbia | ACTTGTAAATACTAAAAGTTGATTTTGC | AGGACATGATCAACGAAGTGG |
| *cam7-2* | AT3G43810 | SALK_074336C | Columbia | TTTCCGAAATACATGCGATAAC | TGTGCAGAGATTCACGATCAC |
| *PP2C76-1* | AT5G53140 | CS66142 | Columbia | GTTGTTGCTGAACCCGAGATACAA | ATCAGTTAAGTTATGCGCGAAGGA |
| *PP2C76-2* | AT5G53140 | SALK_010368C | Columbia | GAATGCCAAATCTTATGCGAG | GCTGAACGAATCAGCTTTTTG |
| *LBa1 (5' to 3'):* TGGTTCACGTAGTGGGCCATCG | | | | | |

**Table S3. Primers for RT-PCR.**

| | |
|---|---|
| Semiquantitative RT-PCR analysis of CRK gene expression in leaves 2 days after infiltration with 10 mM MgCl2 (mock) or *Pst* DC3000 | |
| Names of the primers | Primer sequences (5’ to 3’) |
| CRK12 F | GGTATTACGACTCTACACTTCCAACAAC |
| CRK12 R | TCAACGAGGATCTAGATTTGTAATGG |
| CRK14 F | TTTGTTGTTTATAGGAGGAGGAAATCATAC |
| CRK14 R | TCAACGAGGTTCAAAATCTGTAATGGTTAC |
| CRK16 F | AGGAGGAGAAAAGCCTACCAAGAGTTTG |
| CRK16 R | ACGAAGATCAACACTCGTGATCGATG |
| CRK17 F | TGCAAGAGAAGAAAGCAAAAGCAAG |
| CRK17 R | TCAATTTCTTGTAGAAATCTTTCCGCCATC |
| CRK18 F | AGAAGAAAGCAAAAGCAAGAAATGGATC |
| CRK18 R | TCAACGAGGATTAACGTCCGTGATTG |
| CRK21 F | TAGGAGGAGAAAAGCTTACCAATC |
| CRK21 R | TTAACGAGGTCTAACACTCGTGATC |
| CRK24 F | TGGAAGAGGAGAAAAGCATATAAAACAAAG |
| CRK24 R | ACGAGGACTAACACACGTGACCGATAC |
| CRK30 F | TGCAGGAGTAGAAAAAAATATCAAGC |
| CRK30 R | TCAATCCTCAGTGTTTCTATACATTGAC |
| CRK31 F | AAGCGAAGACAATCATACAAAACACTG |
| CRK31 R | TCACCGAGGAGTGGCTCTAGTGATTG |
| CRK32 F | CGGAAGAGAAGACAATCGTACAAAAC |
| CRK32 R | TTACCGAGGAGTGACTCTAGTGATTG |
| CRK33 F | GTTTGCAGAAAGAGAAAAACTGATC |
| CRK33 R | TCAGCGAGGAACTAAGTCATCAATC |
| Semiquantitative RT-PCR analysis of the gene expression in WT and T-DNA insertion lines | |
| CRK21 F | AGCGTCCTCAGGTTCGCTTCG |
| CRK21 R | TGCTGGCTTTGAGGTCACGGT |
| MKK3 F | GAATTCAATGGCTATGTGTGCTAC |
| MKK3 R | GGTGAGCATATCAGCTAAATCTTTC |

| | |
|---|---|
| PP2C76 F | ATGGTATGCAGCAGTTTCATAAG |
| PP2C76 R | CATGGCCATCAAATATTCCAAAC |
| CaM7 F | ATGGCGGATCAGCTAACC |
| CaM7 R | TCACTTTGCCATCATGACTTTG |
| Actin2 F | ACACTGTGCCAATCTACGAGGGT |
| Actin2 R | GCTGTGATTTCTTTGCTCATACGG |
